# Structure based discovery of antipsychotic-like TAAR1 agonists

**DOI:** 10.1101/2025.10.16.682895

**Authors:** Sijie Huang, Heng Liu, Yujin Wu, Joao M. Braz, Divya Kranthi, Brendan W. Hall, Xinyue Zhang, Dmytro S. Radchenko, Yurii S. Moroz, John J. Irwin, Allan I. Basbaum, H. Eric Xu, William C. Wetsel, Brian K. Shoichet

**Author notes:** These authors contributed equally: Sijie Huang, Heng Liu, Yujin Wu. Corresponding authors (Brian K. Shoichet), (William C. Wetsel), (H. Eric Xu).

## Abstract

Schizophrenia is a severe mental illness whose current treatments primarily target dopamine and serotonin receptors. These drugs often cause side effects and vary in effectiveness across patients. The trace amine-associated receptor 1 (TAAR1), which modulates monoamine signaling, has emerged as a promising alternative target. To discover new TAAR1 ligands, we computationally docked 65 million molecules against the active state of TAAR1 and experimentally tested 55 of those highly ranked. Fourteen molecules active against TAAR1 with potencies ranging from mid-nanomolar to micromolar, all as agonists. This high functional selectivity may reflect the compact conformation adopted by the activated TAAR1 orthosteric site. While this was favorable for agonist prioritization, simulations suggest that it can be over-optimized for initial hit rates at the expense of subsequent affinity maturation. Here, hit optimization yielded nanomolar agonists whose docking-predicted poses were confirmed by cryo-EM. Three agonists had high brain exposure and potencies apparently better than the investigational drug ulotaront and sufficient for behavioral studies. All three potently normalized amphetamine-induced pre-pulse inhibition in mice, a model for schizophrenia, without catalepsy, a common side effect of traditional antipsychotics.

## Introduction

The trace amine-associated receptor 1 (TAAR1) is a G protein-coupled receptor (GPCR) that has received growing attention in recent years for its role in modulating monoaminergic neurotransmission ^1^. TAAR1 is activated by trace amines such as β-phenylethylamine (β-PEA), p-tyramine, tryptamine, and several synthetic psychostimulants, including amphetamine ^2^ and methamphetamine (METH) ^3^; its native neurotransmitter remains uncertain. Unlike other members of the TAAR family, which are predominantly expressed in olfactory regions ^4^, TAAR1 is widely expressed in the brain, particularly in monoaminergic nuclei such as the ventral tegmental area (VTA) and dorsal raphe nucleus (DRN) ^5,6^. This distribution is consistent with its regulation of the dopaminergic ^5^ and serotonergic ^7^ systems, including reward circuits and mood ^8^. These properties make TAAR1 a potential target for psychiatric disorders like schizophrenia, anxiety, depression, and psychostimulant addiction ^6,9–11^. Indeed, the drug SEP-363856 (ulotaront), a potent TAAR1 agonist with joint 5-HT_1A_ activity, was developed as a new therapeutic to treat schizophrenia ^11,12^. Ulotaront was designed to address both the positive and negative symptoms of the disorder while minimizing the side effects often associated with conventional antipsychotics ^13,14^. Despite promising results in early-phase studies, the drug failed to meet the primary endpoints in Phase 3 trials and was not further advanced.

Despite ulotaront’s limitations, TAAR1 remains an interesting target to treat schizophrenia ^15–19^, with new chemotypes being explored including from a successful library docking campaign that targeted a homology model of the receptor^17,20^. Large library docking has recently shown promise against multiple targets^9,21–34^ and here we leverage a newly-determined experimental structure of TAAR1 in its active form, seeking novel agonists. From large library docking emerged agonists with potencies ranging from 23 nM to 20 µM, with an overall hit-rate (number active/number tested) of 25% (14/55). Three agonists were optimized for activity and for pharmacokinetics (PK), reaching TAAR1 potencies from 1 to 48 nM. Encouragingly, they were also active at the 5-HT_1A_ receptor, a co-target with TAAR1 for antipsychotic efficacy^35,36^. While potency was lower against 5HT_1A_, ranging from 260 to 600 nM, these activities were higher than that of ulotaront and likely relevant given the micromolar brain free fractions reached by the molecules. With their high potency and favorable PK properties, three compounds were further evaluated for antipsychotic-like efficacy in mouse models.

## Results

### Large library docking identifies TAAR1 agonists

The ligands were docked against the TAAR1 orthosteric pocket, largely as defined by the configuration of the agonist RO5256390 ^37^ in that site (PDB: 8W8A) (Fig. 1a, b). Using DOCK3.7 ^21,38^, molecule orientations were sampled using pre-defined hot spots (“matching spheres”) as guides. For each orientation, up to 200 ligand conformations were sampled based on precomputed low-energy conformational ensembles ^38,39^. Each ligand configuration was scored electrostatic complementarity using a probe-charge model in precalculated electrostatic potential grids calculated by the Poisson-Boltzmann method QNIFFT ^40^, and for non-polar complementarity using pre-calculated van der Waals terms from the AMBER potential ^41^. Context-dependent ligand desolvation was calculated using SOLVMAP ^42^.

**Fig. 1:**
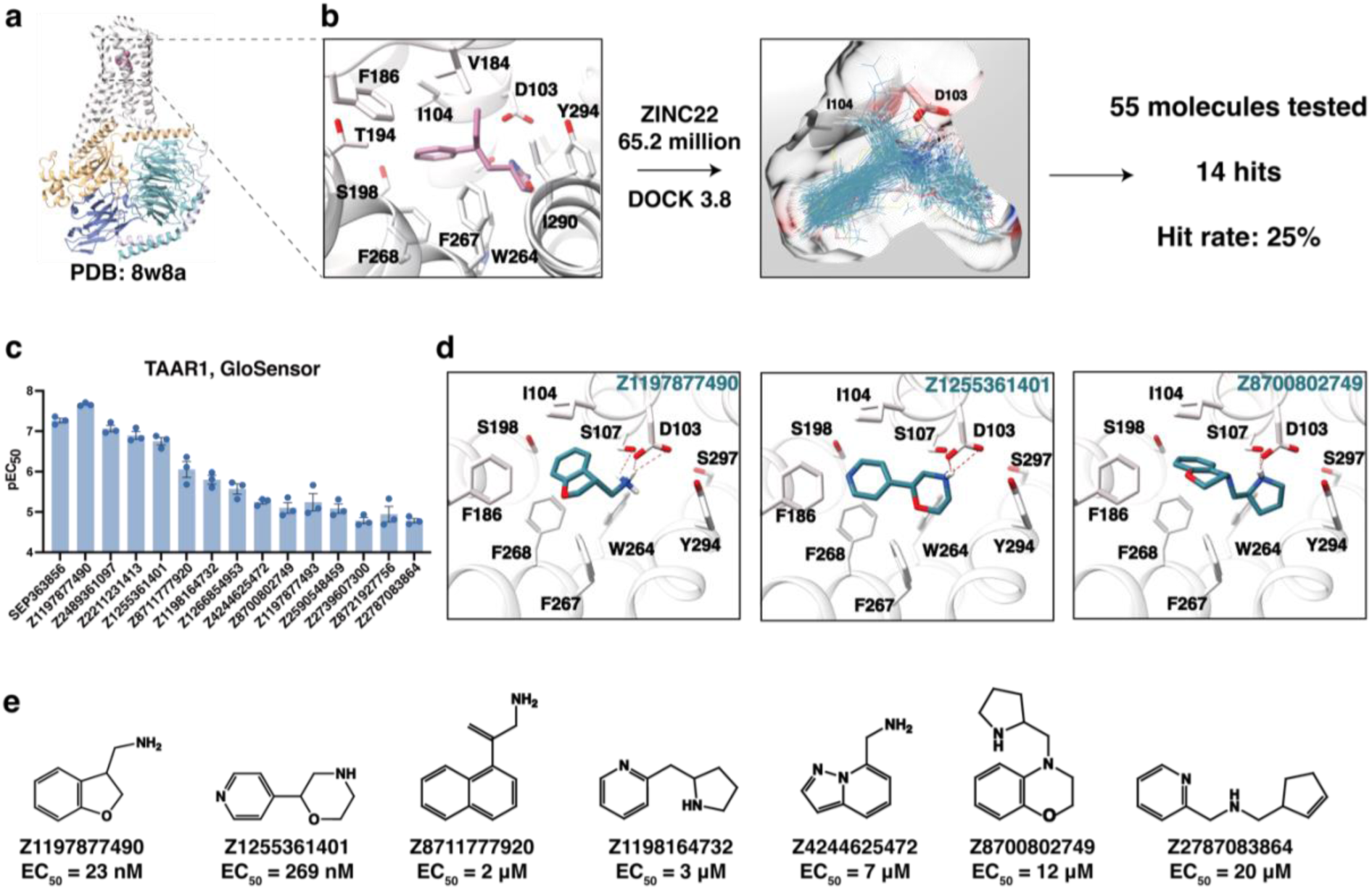
Docking for and *in vitro* testing of TAAR1 agonists. **a,** The cryo-EM structure of the TAAR1 complex (PDB 8w8a). **b,** A library of 65 million compounds was docked to the TAAR1 complex that included G protein. Binding modes of top-ranked compounds from the virtual screening and docked compounds are shown as lines. Hit rates were defined by more than a 30% GloSensor response compared with SEP-363856 at 10 μM. **c,** The GloSensor response of the initial hits at 10 μM. **d,** Docked poses of the hits at the orthosteric site are shown. **e,** Chemical structures of the exemplar hits identified through docking.

As in purely experimental studies, control calculations are important sanity checks before launching prospective docking campaigns^43^. We tested our sampling and the energy potential maps used for scoring by asking whether we could highly-rank known TAAR1 agonists against larger sets of property-matched ^38^ and extrema decoys^43–45^. Electrostatic and desolvation energies may be adjusted by changing the boundaries between low- and high-dielectric in these calculations; we varied these to reach log-weighted AUC values of 31.7 (Extended Data Fig. 1b). This relatively favorable value suggested we could well-distinguish true TAAR1 ligands from decoy molecules.

With these controls in hand, we screened approximately 65 million fragment-like cations (between 11 and 19 non-hydrogen atoms) from the ZINC22 compound library ^46^. This calculation took 39,993 core hours, or less than two days on 1000 cores. Seeking novelty, from the top 382,767 scored molecules we excluded compounds that topologically resembled known TAAR1 agonists; any docked molecule with an ECFP4-based Tanimoto coefficient > 0.35 was removed. To ensure direct interaction between ligand cations and the recognition sites D103^3.32^, Y294^7.43^, F267^6.51^, F268^6.52^, I104^3.33^ and F186^ECL2^ within the orthosteric binding pocket of the receptor, interaction fingerprint-based filters were implemented using LUNA (https://github.com/keiserlab/LUNA). This ensured that the cation was not only well-complemented by the electrostatic potential of the receptor in the area of D103^3.32^—which is what was scored in DOCK3.8—but that a recognizable ion-pair was made. The 95,126 compounds that remained were clustered by topology (see Methods) and the best scoring molecules from the top clusters were visually evaluated using Chimera ^47^. From these, 2,462 diverse molecules, none more similar to one another than an ECFP4 Tc of 0.25, were inspected for favorable geometry and interactions. Sixty-six were prioritized of which 55 were successfully synthesized by Enamine, an 83% fulfillment rate (Fig. 1b).

To evaluate the ability to activate TAAR1, the 55 docking hits were first screened at 10 μM using functional live-cell GloSensor assays ^48^ to evaluate their ability to activate TAAR1 by recruiting Gs and inducing cAMP production (Fig. 1a). Fourteen of the 55 compounds had agonist activity with efficacies (Emax) exceeding 30% of ulotaront’s (SEP363856) (Fig. 1c, Extended Data Fig. 1a and Table 1). Concentration-response experiments confirmed that these compounds acted as TAAR1 agonists, with the negative logarithm of the half-maximal effective concentration (pEC_50_) values ranging from 4.7 to 7.6 and Emax values ranging between 44% and 108% of the ulotaront E_max_ (Fig. 1c and Extended Data Fig. 1c). Based on their potency and novelty (i.e., ECFP4-based Tcs of 0.27, 0.18, and 0.2, respectively), we prioritized compounds Z1197877490, Z1255361401, and Z8700802749 for structure-based optimization (Fig. 1c-e, Extended Data Figs. 2-3 and Table 1–2).

### Only agonists found

In principle, docking against an activated or inactivated state of a receptor should bias toward the discovery of agonists or antagonists, respectively. Whereas we have occasionally observed this to be true for some targets ^25,49–51^, more often we found a mix of agonists and antagonists ^21,28,52–54^. This result is consistent with the idea that docking primarily selects for binding and struggles to distinguish between activating and inactivating ligand interactions. However, in docking against the TAAR1 active state, all 14 actives were agonists and none of the 55 tested were active in antagonist assays (Supplementary Fig. 1). To better understand this result, we asked whether 28 agonist (the 14 new agonists and 14 known agonists) had better fits and ranks when docked to the experimental active-state structure versus a modeled *inactive* structure from Boltz-2 ^55^, comparing also the behavior of seven known antagonists ^7,56,57^. Against the active-state structure of TAAR1, 23 of the 28 agonists ranked among the among the top ∼0.5% of the docked library, with DOCK scores ranging from -54 kcal/mol to -33 kcal/mol. Only one of the seven antagonists fell in this range, with a score of -45 kcal/mol. Docked against the inactive state model, the agonist scores shifted to worse (less negative) values, ranging from -44 kcal/mol to -26 kcal/mol, worsening by an average 8.5 kcal/mol, and their relative ranks suffered even further. Meanwhile, the performance of the antagonists improved, with the best now receiving a score of -46 kcal/mol and two others climbing by between -42 and -35 kcal/mol into the top-ranking molecules (Extended Data Fig. 4).

The unusual ability of the TAAR1 active state docking to prioritize agonists, and of the inactive structure to relatively prioritize antagonists, may reflect the size distinction between agonists and antagonists, and correspondingly between active and inactive receptor conformations. TAAR1 agonists are typically smaller than its antagonists, and the TAAR1 orthosteric site compacts on activation (the site contracts even more in a TAAR1 modeled structure used for docking, a point to which we will return). Another receptor where this is true is the α2A-adrenergic receptor, where docking has also prioritized exclusively agonists from active-state docking ^25^. It may be that docking’s ability to predict ligand function (agonist, antagonist) will be strongest where there are large differences in the physical properties of the active and inactive states—here site volume. Where differences are subtler, docking may struggle to distinguish among ligand functional classes.

### Initial hits optimization

To enhance the potency of the initial docking-active compounds, we employed two strategies. The first involved searching within Enamine REAL Database (e.g., using the SmallWorld program, https://sw.docking.org/) to find similar molecules. In a second strategy, conservative modifications were introduced into to the initial hits wherever synthetically feasible, a strategy that can be remarkably effective ^42^.

We selected 2,012 analogs from the SmallWorld database and explored 27 additional small change derivatives related to Z1197877490. Similarly, for Z1255361401, we retrieved 1,707 analogs from the database and designed 13 new small-change derivatives. For Z8700802749, we identified 873 structurally similar compounds from the database. We docked the analogs to TAAR1 and prioritized a handful for synthesis based on fit (score and visual inspection); five analogs of ‘7490, six of ‘1401, and one analog of ‘2749 were successfully synthesized at Enamine. The parent agonist ‘7490 began with an TAAR1 EC_50_ of 23 nM, and further modifications, such as replacing its oxygen with carbon or sulfur atoms only slightly improved potency (Fig. 2a). Stereochemical purification to EN300-47307925 (Fig. 2a and Extended Data Fig. 2a) also had little effect on TAAR1 potency, though it enhanced 5-HT_1A_ EC_50_ by 26-fold, to 261 nM, nearly ten-fold higher than ulotaront. Conversely, small modifications to the 270 nM ‘1401 substantially enhanced potency (Fig. 2b and Extended Data Fig. 2b). Moving the nitrogen position within the pyridine ring or replacing the pyridine with a benzene increased activity by 59-to 107-fold (Fig. 2b and Extended Data Fig. 2b). Enantiomeric resolution led to Z898713207 (EC_50_ 3 nM) and Z1255382255 (EC_50_ 1 nM), with the latter exhibiting an EC_50_ of 380 nM against 5-HT_1A_ (Fig. 2b and Extended Data Fig. 2b). Finally, for the ‘2749 series, more substantial modifications were necessary to improve potency. Specifically, relocating the cationic nitrogen to a position two carbon atoms away from the benzene ring enhanced activity 37-fold (Fig. 2c and Extended Data Fig. 3a). Enantiomeric separation led to the identification of Z8987197482 with a TAAR1 EC_50_ of 48 nM and 604 nM for 5-HT_1A_ (Fig. 2c and Extended Data Fig. 3a). Docking poses suggested that shortening the linker by one carbon in ‘7482 repositioned the tetralin group closer to the center of the binding pocket, potentially enhancing π-π stacking (Extended Data Fig. 2c).

**Fig. 2:**
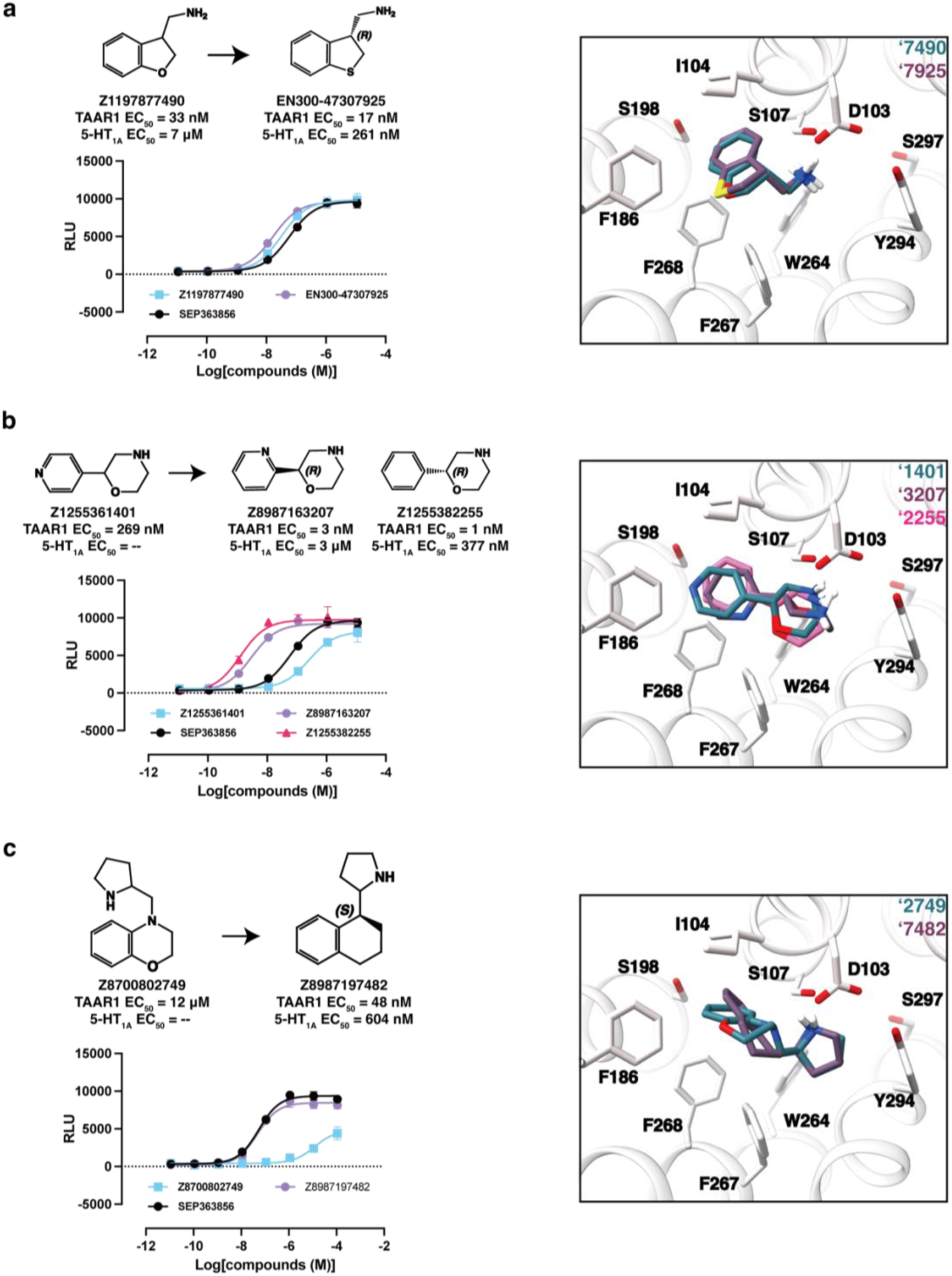
Optimization of ‘7490, ‘1401 and ‘2749. **a,** Docking hit ‘7490 and its optimized analog ‘7925. Superposition of docked poses of ‘7490 and ‘7925 in the orthosteric site. **b,** Docking hit ‘1401 and its optimized analogs ‘3207 and ‘2255. Superposition of docked poses of ‘1401, ‘3207 and ‘2255 in the orthosteric site. **c,** Docking hit ‘2749 and its optimized analog ‘7482. Superposition of docked poses of ‘2749 and ‘7482 in the orthosteric site.

### Structural determination by Cryo-EM

To understand the molecular basis of ligand recognition and provide a template for future optimization, we determined the complex structures of TAAR1 bound to two low-nanomolar agonists: ‘2255, and ‘7482 (Fig. 3a-b, Extended Data Figs. 5-6 and Supplementary data Table 1). The global nominal resolutions of the corresponding cryo-EM density maps for these complexes were 3.14 Å and 3.12 Å, respectively. The overall structures of the complexes resembled previously-reported experimental TAAR1 structures, with root mean square deviation (RMSD) values of 0.492 Å and 0.846 Å, respectively, when compared to the TAAR1 structure 8W8A ^37^.

**Fig. 3:**
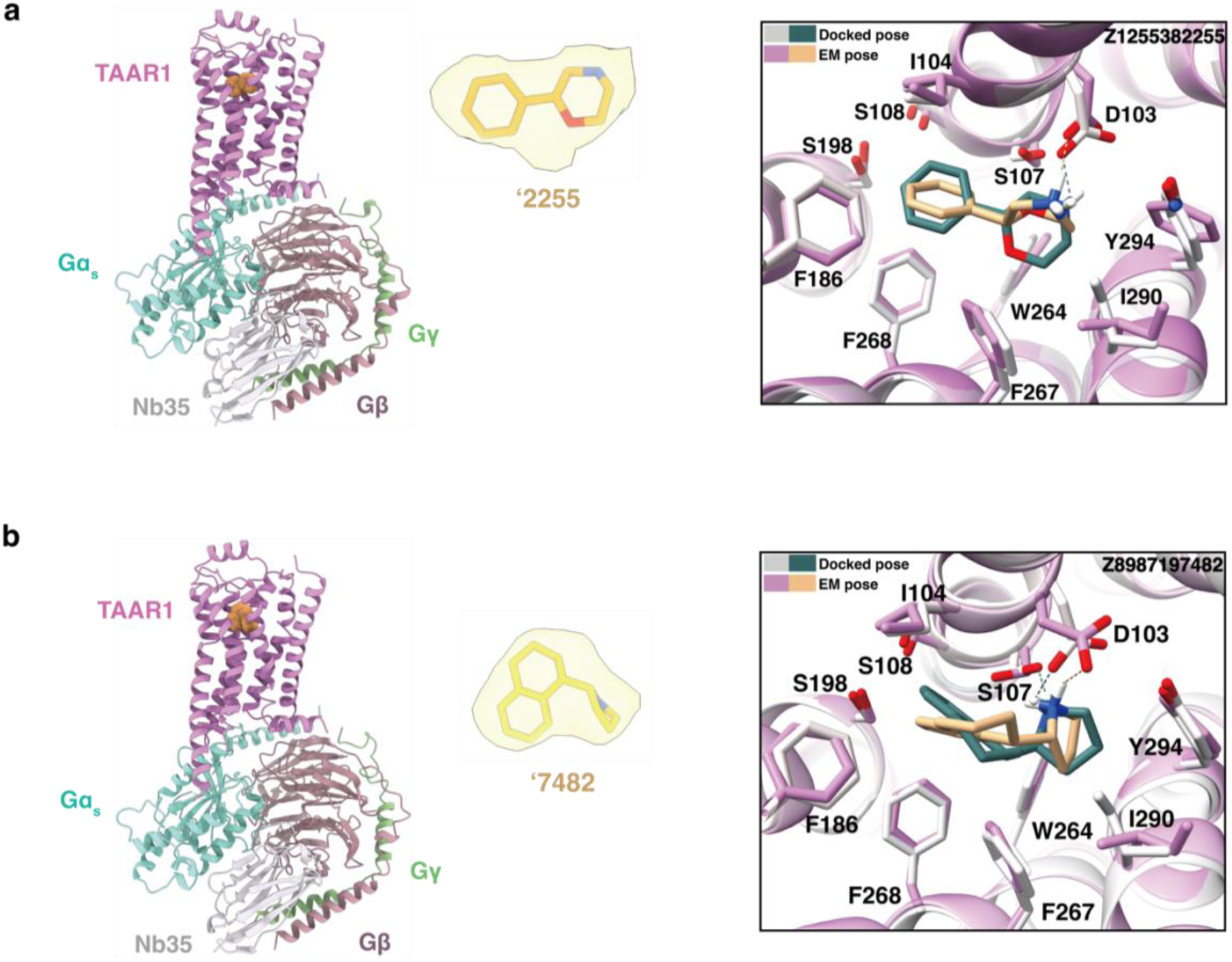
Cryo-EM structures of the ‘2255-TAAR1-Gs and ‘7482-TAAR1-Gs complexes. **a,** Cryo-EM model of ‘2255-TAAR1 highlighting the ligand density. Superposition of the docked and experimentally determined poses of ‘2255 in the orthosteric site. **b,** Cryo-EM model of ‘7482-TAAR1 highlighting the ligand density. Superposition of the docked and experimentally determined poses of ‘7482 in the orthosteric site.

In these two TAAR1 structures, the receptor adopts a canonically activated conformation, with both agonists occupying the orthosteric site and TM6 exhibiting similar degrees of outward displacement. TAAR1 appears to recognize these agonists primarily through two ligand moieties: an aryl group that forms π-π and packing interactions with a sub-pocket formed by F268^6.52^, F186^ECL2^, I104^3.33^, and a cationic amine that ion-pairs with D103^3.32^. These interactions are both predicted by the docking and are captured in the cryo-EM structure. The docked-predicted and cryo-EM observed agonist poses superpose with RMSD values of 2.3 Å and 1.4 Å for ‘2255 and ‘7482, respectively (Fig. 3a-b).

Although the experimental and predicted agonist poses make the same receptor interactions, there were meaningful differences between the structures. In the experimental pose of ‘2255, the ligand undergoes a conformational rotation from its relatively extended and planar docking pose. This reorientation positions the phenyl ring at an inclined angle, facilitating a π-π interaction with F268^6.52^ while at the same time bringing it closer to F186 on ECL2. Similarly, the cationic nitrogen undergoes a shift within the saturated ring system to optimize salt bridge formation with D103^3.32^. As the cationic nitrogen moves closer to D103^3.32^, the residue is slightly displaced outward. Similarly, ‘7482 shifts toward ECL2, bringing its phenyl ring closer to F268^6.52^. Its cationic nitrogen also moves closer to D103^3.32^, resulting in a greater lateral displacement of this residue compared to the ‘2255-bound structure (Fig. 3a-b). More broadly, both the experimental and docking structures of these two compounds help to explain the SAR observed with these two series. For instance, the move from the pyridine of the 3 nM ‘3206 to the phenyl of the 1 nM ‘2255 is consistent with the placement of the phenyl in the non-polar, “aryl cage” of TAAR1. In this geometry, the pyridine would pay a desolvation cost without compensating polar interactions in the site, while the phenyl would not pay this cost. Similarly, the move from the morphilino moiety of ‘2749 (12 µM) to the cyclohexyl moiety of ‘7482 (48 nM), where both are posed in a non-polar sub-pocket, helps to explain the improved potency of the latter (the reduced planarity and better fit of the latter likely also contributes to this >250-fold improvement).

### Comparison to docking against AlphaFold2 models of TAAR1

In a fascinating recent study, library docking was used to find novel agonists targeting an AlphaFold2 (AF2) model of TAAR1 ^20^. Beginning with 100 different models, the authors selected those that best enriched known TAAR1 ligands versus decoys and used these to template a prospective search of a large library, a strategy they had helped pioneer ^58^. On testing of high-ranking molecules, 60% of their predictions were active, with the best of them having an EC_50_ of 35 nM. By hit-rate, these results are substantially better than what we found here using essentially the same docking program, while the affinities are similar for the two screens. Despite the fact that the AF2 model was in an inactivate conformation (Fig. 4a), all of their new ligands were functionally agonists. How may these observations be reconciled?

**Fig. 4:**
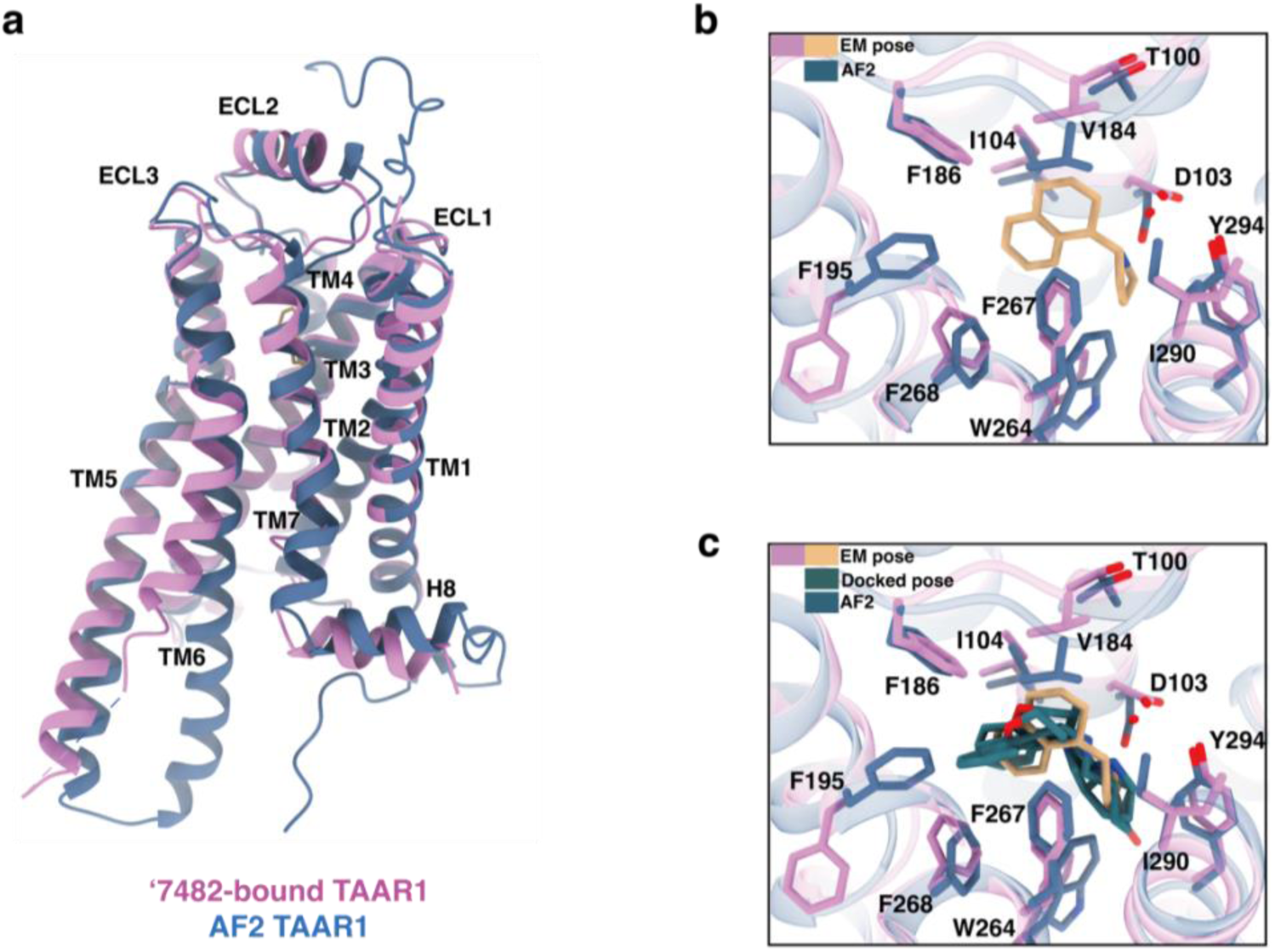
Comparison of the experimental TAAR1 structure and an AlphaFold TAAR1 model^20^, along with the positions of docking-predicted molecules in both structures. **a,** Superposition of ‘7482-bound experimetal TAAR1 and AlphaFold TAAR1 structure. **b,** Superposition of the experimentally determined pose of compound ‘7482 with the AlphaFold TAAR1 structure in the orthosteric site. **c,** Superposition of the experimental ‘7482-bound TAAR1 orthosteric site with several docking hit poses and the orthosteric site of the AlphaFold TAAR1 model.

Ordinarily one might expect an experimental structure, especially one that adopts the full activated state, as our cryoEM structure does, to perform at least as well as an AF2 model, and to better bias toward agonists. One advantage of the AF2 approach is that multiple models were calculated and those that best-enriched ligands in control calculations were used prospectively. While this likely contributed to the high prospective hit rate, it may also have had unanticipated consequences. One reason that the AF2 model may have been so successful is that it represents an unusually tight orthosteric site, with multiple residues clamping down towards the ligands site versus the experimental structure (Fig. 4b), leading to a binding site whose volume ranges from 513 to 639 Å^3^, depending on the model, substantially smaller than the 746 Å^3^ of the experimental structure used here. This will bias the docking toward the small agonist cations favored by TAAR1. But the AF2 orthosteric site was closed down to such an extent that other plausible ligands, including many of those found in our study, could no longer fit (Fig. 4c). It may have also made it harder to optimize the initial molecules for potency because the ligands had few places to grow. Thus, the potency of the best agonist from the AF2 docking, after optimization, was little changed from the initial docking hit. Meanwhile, the larger orthosteric site of the cryoEM structure allowed us to optimize into the 1 nM range, something that ultimately contributed to their high in vivo potency (below). It may be that the strategy of selecting model structures by ligand enrichment— something widely used in the field, including by ourselves ^59–61^—may optimize for initial discovery but lead to challenges for optimization.

### In vivo antipsychotic-like activity of the novel TAAR1 agonists

To assess the therapeutic potential of compounds ‘3207, ‘2255, and ‘7482, we first measured their pharmacokinetics on 10 mg/kg intraperitoneal (i.p.) injection in mice (Extended Fig. 7 and Supplementary data Table 2). All three had high exposure in both brain and cerebrospinal fluid (CSF), the latter of which is a widely used proxy for fraction unbound (Fu) in the brain. Among them, ‘3207 reached a CSF concentration (C_max_) of 3040 ng/mL (about 18 μM), approximately twice that of ‘2255 and ‘7482. Additionally, the area under the concentration-time curve (AUC) in the CSF was 292,000 ng·min/mL for ‘3207 versus 67,400 ng·min/mL and 123,000 ng·min/mL for ‘2255 and ‘7482, respectively. The half-life (T_1/2_) of ‘3207 in CSF was 56 minutes, whereas ‘2255 and ‘7482 had T_1/2_ values of 35 and 64 minutes, respectively. Collectively, ‘3207, ‘2255, and ‘7482 combine high potency at TAAR1 with high brain exposure, leading to receptor coverage (free concentration/EC_50_) between 1000 to 10,000-fold. We therefore evaluated their antipsychotic-like efficacies in pre-pulse inhibition (PPI), a behavioral test that is well-known to be affected by TAAR1 agonists ^12,62,63^.

PPI of the acoustic startle reflex assesses the ability of a weak pre-stimulus to suppress the response to a subsequent strong startle. PPI deficits are observed in patients with schizophrenia ^64^, and similar disruptions in rodents are used to model psychotic-like states ^65^. To induce such deficits, we administered (i.p.) amphetamine (AMPH) to mice—a well-established method for disrupting PPI in rodents. These deficits are known to be reversible by antipsychotic drugs.

Mice were injected (i.p.) either with vehicle (Veh) or with the new TAAR1 agonists (Fig. 5a-c). Ten minutes later, they were given Veh or 3 mg/kg AMPH. Thus, the treatments consisted of a Veh-Veh control, a compound-Veh control, a Veh-AMPH positive control, and the compound-AMPH test group (see Supplementary Table 3 for statistics; *post-hoc* values are displayed in the figures). Results from the ‘3207 experiment revealed PPI was disrupted in the Veh-AMPH group relative to the Veh-Veh control and the 0.5 mg/kg ‘3207-Veh animals (Fig. 5a). This AMPH-induced PPI impairment was normalized with 0.5 mg/kg ‘3207. Similarly, in the ‘2255 experiment the Veh-AMPH treatment depressed PPI *versus* the Veh-Veh, the 0.2 mg/kg ‘2255-Veh, and the 0.2 mg/kg ‘2255-AMPH groups (Fig. 5b). PPI in the 0.1 mg/kg ‘2255-AMPH mice was reduced versus the Veh-Veh and 0.2 mg/kg ‘2255-AMPH groups, indicating that the 0.2 mg/kg dose of ‘2255 more effectively reversed the AMPH-induced PPI disruption. Finally, tests with ‘7482 demonstrated the Veh-AMPH treatment impaired PPI *versus* the Veh-Veh and 0.2 mg/kg ‘7482-AMPH groups (*p*-values≤0.050), indicating that 0.2 mg/kg ‘7482 effectively restored the PPI deficit. (Fig. 5c). Collectively, the ‘3207, ‘2255, and ‘7482 normalized the Veh-AMPH induced disruption in PPI, with the ‘2255 and ‘7482 being more efficacious than ‘3207, likely reflecting their higher potency on target. All three compounds potently reduced PPI in the AMPH mouse models.

**Fig. 5:**
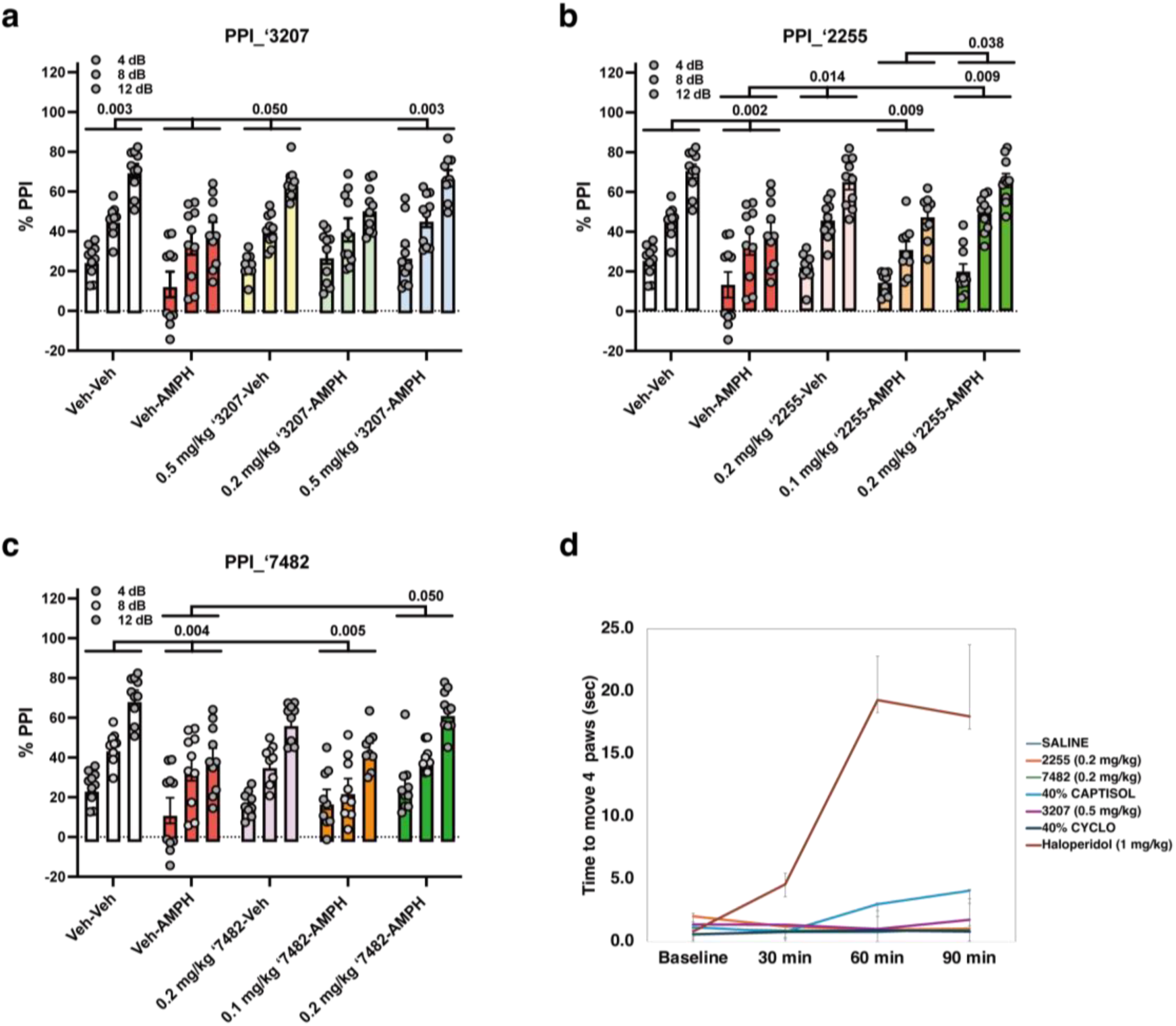
Effects of ‘3207, ‘2255, and ‘7482 on restoring amphetamine-disrupted pre-pulse inhibition. **a-d,** Mice were administered the vehicle (Veh) or the new agonists, followed 15 min later with the Veh or 3 mg/kg amphetamine (AMPH). They were placed in the PPI apparatus and tested after given 10 min of habituation. Effects on PPI of administration **(a)** ‘3207 **(b)** ‘2255 **(c)** ‘7482. **d,** In the catalepsy test doses of the TAAR1 agonists ‘2255, ‘7482 and ‘3207 that rescued AMPH-disrupted PPI, did not induce catalepsy. Conversely and as expected, the positive control haloperidol induced catalepsy in mice that lasted at least 90 minutes post-injection. The data are presented as means and standard errors of the mean. In each a-c panel the *post-hoc p*-value is provided for a specific comparison. The primary statistics are in Supplementary Table 3.

Besides PPI, null and startle activities and effects on catalepsy were also evaluated for the three agonists. Null activity was consistently low in Veh-Veh and compound-Veh groups compared to Veh-AMPH and at least one compound-AMPH group (Supplementary Fig. 2a-c). For pulse or startle activities, the patterns of responses differed slightly among the three compounds but overall, pulse activities were lower in the highest-dose groups of each compound compared to other groups (Supplementary Fig. 2d-f). We also evaluated the potential for inducing catalepsy, a serious adverse drug reaction of many current antipsychotics. At doses that rescue amphetamine-disrupted PPI, mice treated with ‘2255 (0.2 mg/kg i.p.), ‘7482 (0.2 mg/kg i.p.), or ‘3207 (0.5 mg/kg i.p.) showed no observable catalepsy at 60 or 90 min, unlike those treated with the classic dopamine receptor antagonist haloperidol (1 mg/kg i.p.) (Fig. 5d). This suggests that the new agonists have a favorable catalepsy-related safety profile versus classic anti-dopaminergics. Based on literature values for pre-pulse inhibition in mice, the new molecules, with substantial activity at doses as low as 0.2 mg/kg, are at least as potent and arguably more potent than drugs like ulotaront.

## Discussion

Four observations from this study merit emphasis. **First**, library docking and structure-guided optimization found agonists with nanomolar potency on TAAR1 and with sub-µM activity on 5-HT1A, making them more potent both in vitro and apparently in the mouse than ulotaront. **Second,** only agonists were found. This may reflect the choice of an active-state structure of the TAAR1 for docking, where the orthosteric site contracts around the agonist, selecting for smaller agonist-like molecules and discriminating against larger antagonist-like molecules. Consistent with this view, a previous docking campaign versus AlphaFold2 structures of TAAR1^20^, where the orthosteric site contracted even more, had even higher agonist hit-rates, even though the overall TAAR1 model was in an inactive state. While the compact orthosteric site of the AlphaFold2 model may have improved hit-rates, it may also have restricted the ability to optimize for more potent leads because it was overly compact, something perhaps selected for in optimizing the docking parameters against the AlphaFold2 structure. **Third**, the docking predictions here were largely confirmed by the subsequent cryo-EM structures, which may template further optimization of these compounds. **Fourth,** three of the new agonists potently restored AMPH-disrupted PPI without the catalepsy that is a key liability for many of the classic dopamine receptor antagonists. These new TAAR1 agonists may possess a broader therapeutic index than typical approved antipsychotic drugs for treating acute psychosis.

Several caveats bear mentioning. Although we found agonists with improved potency for both TAAR1 and 5-HT_1A_ receptors, their primary target remains TAAR1, with only moderate potency against 5-HT_1A_. While the new agonists are topologically dissimilar to known TAAR1 ligands, they share physical features—and in some cases, core scaffold elements—likely due to the receptor’s small orthosteric site. Finally, the in vivo activity of ‘3207, ‘2255, and ‘7482 were only evaluated in the PPI model of AMPH-disruption for acute psychosis. Their potential effects on other behavioral phenotypes, as well as other adverse drug reactions, remain unexplored.

These limitations should not obscure the core findings from this study. Large library docking against TAAR1 followed by structure-based optimization revealed a series of low-nanomolar TAAR1 agonists with topologies unrelated to known ligands. Targeting the active state of the receptor successfully biased toward agonist discovery. Cryo-EM structures largely supported the docking predictions. The favorable physical properties of the new TAAR1 agonists, something insisted upon in the choice of library and docking hits, led to molecules with high CNS exposure and behavioral changes consistent both with improving sensorimotor gating and reducing the side effects of catalepsy. With structures and pharmacology differing from known investigational drugs against TAAR1, these results support the further optimization of these new agonists and, more broadly, a structure-based approach to ligand discovery for this interesting receptor ^17,20^.

## Materials and Methods

### Molecular docking against TAAR1

In the docking, we targeted the active-state structure of TAAR1 represented by the cryo-EM structure in PDB 8w8a. The receptor was protonated assuming a pH of 7.4 using Epik and PROPKA in 2022-released Maestro. Energy grids for the different terms of the DOCK3.8 scoring function were pre-generated—the van der Waals term based on the AMBER force fields using CHEMGRID ^66^, Poisson-Boltzmann–based electrostatic potentials using QNIFFT ^40^, and the context-dependent ligand desolvation using SOLVMAP ^38^. The resulting grids were evaluated for their ability to enrich known TAAR1 ligands over property-matched decoys. We chose 14 known TAAR1 agonists and used the DUDE-Z pipeline ^44^ to generate 700 decoys. After optimization of the dielectric boundaries for the Poisson-Boltzmann electrostatic and the GB/SA desolvation grids, adjusted logAUC values of 34.1 were achieved. We also docked against an “extrema” set from the DUDE-Z web server (http://tldr.docking.org) to ensure that molecules with extreme physical properties were not enriched ^43,44^. We found close to 80% monocations were enriched among the top 1000 ranking molecules, with a logAUC of 15.4. Subsequently, 65 million “fragment-like” molecules (molecular weights ≤ 250 Da and clogP ≤ 3.5) from the ZINC22 database (https://cartblanche22.docking.org) were docked against TAAR1 using DOCK3.8. On average, 19,140 orientations and 408 conformations were sampled per molecule demanding 39,993 core hours or less than two days on 1000 cores.

Once the screen was completed, the 382,767 molecules with scores ≤−33 kcal/mol were filtered for key interactions with D103, Y294, F267, F268, I104 and F186 and for dissimilarity to known ligands; no molecules with Tanimoto coefficients (Tcs) >0.35 to known TAAR1 ligands in ChEMBL, using ECFP4 fingerprints, were accepted. Then the remaining 95,128 molecules were clustered by LUNA 1024-length binary fingerprints with a Tc = 0.25, resulting in 2462 clusters. From these 66 compounds were chosen by human inspection and prioritized for synthesis, of which 55 compounds were successfully made at Enamine.

### Hit optimization

Potential analogs of the agonists emerging from docking were sought in SmallWorld (NextMove Software, Cambridge UK) searches of a 48 billion make-on-demand libraries ^67^. The resulting analogs were docked to the TAAR1 and evaluated for fit. Compounds were also designed by exploring single atom changes to the parent agonists that were synthetically accessible at Enamine. Compound purities were at least 90% and most active compounds were at least 95% (assessed by LC/MS and NMR).

### GloSensor cAMP Accumulation Assay

The Promega GloSensor cAMP Assay ^48^ was performed to monitor the real-time cAMP variation induced by the TAAR1 or 5-HT_1A_R. Full-length TAAR1 and 5-HT_1A_R were cloned into a pcDNA3.1 vector and transfected into HEK293T cells. Before transfection, the cells were cultured in DMEM media (Gibco) with 10% FBS (Thermo Scientific) at ∼70% confluency in six-well plates. After 16 h, cells were transiently transfected following the Lipofectamine 3000 Reagent transfection system (Thermo Fisher) with 0.5 μg receptor and 1.5 μg GloSensor-22F (Promega). After 24 h, transfected cells were digested and transferred into a 96-well plate with warmed CO_2_–Independent media (Gibco) containing 100 μM Beetle luciferin (Promega) for 1 h. One μL of 100x test compounds at various concentrations were added and incubated for 10 min at room temperature. For the 5-HT_1A_R, an additional 10 μL Forskolin was added to the cells at a final concentration of 5 μM. Luminescence measurements were recorded on a CLARIOstar plate reader and monitored for 12 minutes. All observations were analyzed with GraphPad Prism v.10. Non-linear curve fits were performed using a three-parameter logistic equation (log (agonist versus response)).

### Expression, complex formation and purification

The TAAR1 complexes with the new agonists were expressed in Hi5 insect cells (Invitrogen). Cells were cultured in SIM HF Expression Medium (Sino Biological, MHF1) to a density of 3×10^6^ per mL with virus preparations for TAAR1-miniGα_s_ and Gβ_1_γ_2_ at a ratio of 1:1.2. The infected cells were cultured at 27°C for 48 hours before collection by centrifugation and the cell pellets were stored at -80°C for future use.

Protein purification proceeded as previously described ^37^. Briefly, cell pellets from 2 L culture were thawed at room temperature and resuspended in low salt buffer containing 20 mM HEPES pH 7.4, 100 mM NaCl, 5 mM CaCl_2_, 5 mM MgCl_2_, 10% glycerol, protease inhibitor cocktail (Thermo Fisher). Cell membranes were collected by ultra-centrifugation at 30000 rpm for 30 min. The membranes were resuspended and then incubated with 1 μM Z1255382255 or 5 μM Z8987197482, and 20 mU mL^-1^ apyrase (NEB) at 4 °C for 1 hour. The protein was extracted from the membrane by 20 mM HEPES, pH 7.4, 100 mM NaCl, 5 mM CaCl_2_, 5 mM MgCl_2_, 10% glycerol, 1%(w/v) lauryl maltose neopentyl glycol (LMNG, Anatrace), 0.2% (w/v) cholesteryl hemisuccinate Tris salt (CHS, Anatrace) and stirred for 2.5 hour at 4℃. The supernatant was isolated by centrifugation at 30000 rpm for 30 min and then incubated overnight at 4 °C with pre-equilibrated TALON IMAC resin (Clontech). After batch binding, the TALON IMAC resin was washed with 10 column volumes of 20 mM HEPES, pH 7.4, 100 mM NaCl, 5 mM CaCl2, 5 mM MgCl2, 30 mM imidazole, 10% glycerol, 0.1% LMNG (w/v), 0.02% CHS (w/v), 1 μM Z1255382255 or 5 μM Z8987197482 and eluted with the same buffer plus 300 mM imidazole. The eluted protein was incubated with 20μg/ml of Nb35 at 4 °C for another 2 h. The mixture was then purified by SEC using a Superose 6 10/300 GL column (GE healthcare) in 20 mM HEPES, pH 7.4, 100 mM NaCl, 0.00075% (w/v) LMNG, 0.00025% (w/v) CHS and 100 nM Z1255382255 or 500 nM Z8987197482. The fractions of monomeric complex were collected and concentrated to 5.5-8.0 mg mL^-1^ for electron microscopy experiments.

### Cryo-EM data collection and processing

For cryo-EM grid preparation, samples for Z1255382255 or Z8987197482 were applied individually to EM grids (Quantifoil, 300 mesh Au R1.2/1.3) that were glow-discharged for 45s and then blotted for 4s under 100% humidity at 4 °C before being plunged into liquid ethane cooled by liquid nitrogen using a Mark IV Vitrobot (FEI). Cryo-EM imaging was collected on a Titan Krios equipped with a Falcon 4 direct electron detection device at 300 kV at the Shanghai Advanced Electron Microscope Center, Shanghai Institute of Material Medica. Images were taken with a pixel size of 0.73 Å, a defocus ranging from -1.0 to -2.0 μm using the EPU software (FEI Eindhoven, Netherlands). Each micrograph was dose-fractionated to 36 frames with a total dose of 50 e^-^ Å^−2^ on each EER format movie.

All dose-fractioned images were motion-corrected and dose-weighted by MotionCorr2 ^68^ software, and their contrast transfer functions were estimated by patch CTF estimation in cryoSPARC v3.2.0 ^69^. For the Z8987197482-TAAR1-Gα_s_ complex, particle selection and 2D, 3D classifications were performed using cryoSPARC. The auto-picking process and Topaz extract produced 1,344,129 particles, which were subjected to interactively 2D and 3D classifications, producing a well-defined subset with 137,014 particles. Subsequent non-uniform refinement and local refinement generated a map with an indicated global resolution of 3.12 Å. For the Z1255382255-TAAR1-Gα_s_ complex, particle selection and 2D, 3D classifications were performed using cryoSPARC v3.2.0. The Topaz extract process produced 853,711 particles, which were subjected to interactively 2D and 3D classifications, producing 72,449 particles with complete complex and good quality. The particles were further subjected to non-uniform refinement and local refinement generated a map with an indicated global resolution of 3.14 Å.

### Model building and refinement

The active state of TAAR1 from a previous structure (PDB ID: 8W8A) was used as the starting model for model building the TAAR-Gα_s_ complexes. Restraints for Z1255382255 or Z8987197482 were generated using Phenix elbow. The cryo-EM model was docked into the electron microscopy density map using Chimera-1.14 ^70^, followed by iterative manual adjustment and rebuilding in COOT-0.9.6 ^71^ and ISOLDE-1.2 ^72^. Restraints for Z1255382255 or Z8987197482 were generated using Phenix.elbow ^73^. Real space and reciprocal space refinements were performed using Phenix programs with secondary structure and geometry restraints. The final refinement statistics are provided in Supplementary Table S3. UCSF Chimera-1.14, Chimera X ^74^, and PyMOL-2.0 (https://pymol.org/2/) were used to prepare the structural figures in the paper.

### Pharmacokinetics

The pharmacokinetics (PK) of ‘3207, ‘2255, and ‘7482 were investigated by Bienta Enamine Biology Services (Kiev, Ukraine) in accordance with the study protocols P081724a, P100824a, and P100824b. Compound ‘3207 was formulated in Captisol– Water for injection (40:60, w/v), while compounds ‘2255, and ‘7482 were formulated in saline. For all three studies, a single 10 mg/kg intraperitoneal (i.p.) dose was administered and samples collected at nine time points: 5, 15, 30, 60, 120, 240, 360, 480, and 1440 minutes. Each of the time-point treatment groups included 3 animals. Mice were injected (i.p.) with 2,2,2-tribromoethanol at the dose of 150 mg/kg prior to drawing the blood. CSF was collected under a stereomicroscope from *cisterna magna* using 1 mL syringes. Blood collection was performed from the orbital sinus in microtainers containing K3EDTA. Animals were euthanized by cervical dislocation after the blood samples collection. Each blood sample was centrifuged for 10 min at 3000 rpm. Brain samples were collected and weighed. The samples were immediately processed, flash-frozen, and stored at -70°C until subsequent analysis. The concentrations of the test compound below the lower limit of quantitation (LLOQ = 5 ng/ml for plasma; 5 ng/g for brain, and 10 ng/ml for CSF samples) were designated as zero. Results were analyzed using noncompartmental, bolus injection or extravascular input analysis models in WinNonlin 5.2 (PharSight). Data below the LLOQ were presented as missing values to improve the validity of T_1⁄2_ calculations. For each treatment condition, the final concentration values at each time-point were analyzed for outliers using Grubbs’ test with the level of significance set at p < 0.05.

### In Vivo Behavioral Studies

Adult male and female C57BL/6J mice (#000664; Jackson Laboratories, Bar Harbor, ME) were used. Mice were housed at 3-4 per cage in a temperature- and humidity-controlled room with a 14:10 hr (lights on at 0600 hours) light-dark cycle and provided with food and water *ad libitum*. All PPI studies were conducted with an approved protocol from the Duke University Institutional Animal Care and Use Committee and all investigations and procedures were performed in accordance with ARRIVE guidelines.

The drug consisted of *D*-amphetamine hemisulfate salt (AMPH, #A5880; Sigma-Aldrich, St. Louis, MO) and the compounds included Z8987163207 (‘3207), Z1255382255 (‘2255), and Z8987197482 (‘7482). In the experiments, the groups consisted of two separate administrations (i.p.) separated by 10 minutes consisting of the vehicle (Veh) or different doses of one of the compounds followed with the Veh or 3 mg/kg AMPH. The injection series consisted of the Veh-Veh control, the “compound”-Veh control, the Veh-AMPH positive control, and the “compound”-AMPH test group. The first Veh consisted of *N,N*-dimethyllacetamide (final volume 0.5%; Sigma-Aldrich) that was brought to volume with 5% 2-hydroxypropoyl-β-cyclodextrin (Sigma-Aldrich) in water (Mediatech Inc., Manassas, VA), while the second Veh was water. The AMPH was reconstituted in water. All drugs were administered (i.p.) in a 5 ml/kg volume.

### Prepulse inhibition (PPI)

PPI of the acoustic startle response was conducted using SR-LAB chambers (San Diego Instruments, San Diego, CA) as described ^75^. Mice were injected with Veh; 0.1 or 0.2 Z2255 or Z7482; or 0.2 or 0.5 mg/kg Z3207 and returned to their home-cages. Ten minutes later the mice were given the Veh or 3 mg/kg AMPH and were placed into a Plexiglas holding tube and put in the apparatus. After 10 min of habituation to a white noise background (64 dB), testing began. A test session consisted of 42 trials with 6 null trials, 18 pulse-alone trials, and 18 prepulse-pulse trials. Null trials were composed of exposure to the white noise background, pulse trials contained of 40 msec bursts of 120 dB white-noise, and prepulse-pulse trials were composed of 20 msec pre-pulse stimuli that were 4, 8, or 12 dB above the white-noise background (6 trials/dB), and were followed by the 120 dB pulse stimulus 100 msec later. Testing began with 10 pulse-alone trials followed by combinations of the prepulse-pulse and null trials and was terminated with 10 pulse-alone trials. PPI responses were calculated as %PPI = [1–(pre-pulse trials/startle-only trials)]*100.

All statistical analyses were performed with IBM SPSS Statistics 29 programs (IBM, Chicago, IL). The data are displayed as means and standard errors of the mean. As no sex effects were detected in any experiments, this variable was collapsed. Note, to reduce the numbers of mice used, the same raw data for the Veh-Veh and the Veh-AMPH groups were used within comparisons with the Z3207, Z2255, and Z7482 treatments. The PPI data were analyzed with repeated-measures ANOVA (RMANOVA), with Greenhouse-Geisser corrections (due to violation of homogeneity of variance), followed by Bonferroni corrected pair-wise comparisons. The null and pulse activities were analyzed by one-way ANOVA followed with Bonferroni corrections. If Levene’s test for homogeneity of variance was violated, the data were analyzed by one-way Kruskal-Wallis tests followed with Dunn *post-hoc* tests. A *p*<0.05 was considered significant. The statistics are in Table S5, and all results were plotted using GraphPad Prism 10.4.1.

### Catalepsy Test

Thirty minutes after an i.p. injection of the compounds, mice were placed on a vertical wire mesh and the latency to move all four paws was recorded ^76^.

## Acknowledgements

Supported by internal UCSF funds (to BKS). The cryo-EM data were collected at Advanced Center for Electron Microscopy at Shanghai Institute of Materia Medica, Chinese Academy of Sciences. We are grateful to Kai Wu and Wen Hu for collecting the cryo-EM data. This work was supported by National Natural Science Foundation of China (32130022 and 82121005 to H.E.X., 32501066 to H.L.); the National Key R&D Program of China (2022YFC2703105 to H.E.X., 2019YFA0904200); CAS Strategic Priority Research Program (XDB37030103 to H.E.X.); Shanghai Municipal Science and Technology Major Project (2019SHZDZX02 to H.E.X.); the Lingang Laboratory (LG-GG-202204-01 to H.E.X.); State Key Laboratory of Drug Research (SKLDR-2023-TT-04 to H.E.X.), the Natural Science Foundation of Shanghai (25ZR1402552 to H.L.).

## Author contributions

S.H. and B.K.S. designed the project, with input from H.L., H.E.X., W.C.W. and Y.W. S.H. Y.W. D.K. and B.W.H performed the docking screens with input from B.K.S. Ligand optimization was performed by S.H. with input from B.K.S. H.L. and X.Z. prepared the protein samples for cryo-EM data collection, structure determination, and modelled the structure with input from H.E.X. Y.S.M. and D.S.R. supervised synthesis of compounds at Enamine. S.H. performed GloSensor cAMP assay. Mr. Christopher Means and Ms. Ann Njoroge conducted the PPI studies at Duke and Dr. Ramona M. Rodriguiz analyzed the PPI data as overseen by W.C.W. J.M.B. performed and analyzed catalepsy test results, supervised by A.I.B. The manuscript was drafted by S.H. and edited by W.C.W. and B.K.S., who also supervised the docking and designed many parts of the project. All authors reviewed and approved the final manuscript.

## Data availability

The ZINC compound library is available to all at https://cartblanche22.docking.org/. The PDB entry for the TAAR1 cryo-EM structure used for docking calculations is 8W8A. The ‘2255-TAAR1-Gs model was deposited to the Protein Data Bank under accession code 9VZ3 and the map was deposited to the EMDB under accession code EMD-65477. The ‘7482-TAAR1-Gs model was deposited to the Protein Data Bank under accession code 9VZ2 the map was deposited to the EMDB under accession code EMD-65476. A list of all de novo compounds and their 2D structure can be found in Table S1 and S2. All compounds may be ordered from Enamine.

## Code availability

DOCK3.8 has been previously published and is available to all academics without charge (https://dock.compbio.ucsf.edu/DOCK3.8/).

## Data and Materials Availability

All data needed to evaluate the conclusions in the paper are present in the paper and/or the Supplementary Materials or is available upon request.

## General considerations

All chemicals for the synthesis of the studied compounds were provided by Enamine Ltd. (www.enamine.net). All solvents were treated according to standard methods. ^1^H NMR spectra were recorded at 400 or 500 MHz (Varian or Bruker spectrometers). ^1^H chemical shifts are calibrated using residual nondeuterated solvent DMSO or CHCl_3_: δ = 2.50 ppm or 7.29 ppm, respectively. Coupling constants are given in Hz. LC/MS analysis was performed utilizing Agilent 1200 Series LC/MSD system with DAD/ELSD (column Zorbax SB-C18 1.8 µm 4.6×15mm; solvent A (water, 0.1% formic acid) and solvent B (acetonitrile, 0.1% formic acid); gradient 0% – 100% solvent B, run time, 1.8 min; flow rate, 3 mL/min) and Agilent LC/MSD SL (G6130A), SL (G6140A) mass-spectrometer (APCI mode). All the LC/MS data were obtained using positive/negative mode switching.

## Spectral description for the studied compounds

**1-(2,3-dihydro-1-benzofuran-3-yl)methanamine – ZINCbh0000002DSB, Z1197877490.**

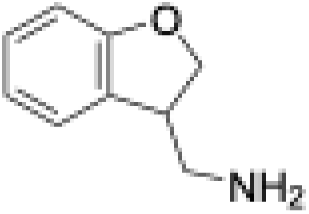

Purity, >95% (assessed by LC/MS).

^1^H NMR (400 MHzCDCl_3_) δ 7.23 (d, *J* = 7.3 Hz, 1H), 7.16 (t, *J* = 7.8 Hz, 1H), 6.88 (t, *J* = 7.4 Hz, 1H), 6.82 (d, *J* = 8.0 Hz, 1H), 4.66 (t, *J* = 9.0 Hz, 1H), 4.43 (dd, *J* = 9.0, 5.2 Hz, 1H), 3.51 (p, *J* = 6.1 Hz, 1H), 2.97 (qd, *J* = 12.5, 5.9 Hz, 2H), 1.24 (s, 2H).

LC/MS (APSI) m/z [M] calculated for C9H11NO: 149.0; found: 149.1.

**1-{4H,6H,7H-thieno[3,2-c]pyran-4-yl}ethan-1-amine hydrochloride – ZINCcn0000002ry9, Z2211231413.**

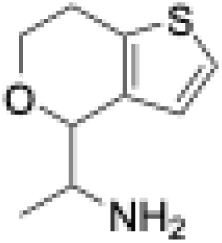

Purity, >95% (assessed by LC/MS).

^1^H NMR (400 MHz, DMSO-*d*_6_) δ 8.11 (d, *J* = 183.0 Hz, 3H), 7.40 (d, *J* = 5.2 Hz, 1H), 6.99 (d, *J* = 5.3 Hz, 1H), 4.85 (d, *J* = 103.7 Hz, 1H), 4.36 – 4.04 (m, 1H), 3.90 – 3.59 (m, 2H), 2.91 (d, *J* = 17.3 Hz, 1H), 2.77 (t, *J* = 15.2 Hz, 1H), 1.13 (dd, *J* = 179.8, 6.6 Hz, 3H). LC/MS (APSI) m/z [M+H] calculated for C9H14NOS: 184.1; found: 184.1.

**1-{5H,6H,7H-cyclopenta[b]pyridin-7-yl}methanamine dihydrochloride – ZINCbg0000002DEe, Z2489361097.**

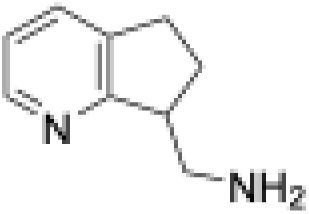

Purity, >95% (assessed by LC/MS).

^1^H NMR (400 MHz, DMSO-*d*_6_) δ 8.59 (d, *J* = 5.5 Hz, 1H), 8.48 (s, 3H), 8.24 (d, *J* = 7.7 Hz, 1H), 7.70 (dd, *J* = 7.7, 5.6 Hz, 1H), 3.85 (p, *J* = 7.4 Hz, 1H), 3.56 (dt, *J* = 11.9, 5.7 Hz, 1H), 3.07 (dddd, *J* = 44.2, 24.8, 15.6, 7.3 Hz, 3H), 2.42 (dtd, *J* = 13.6, 8.6, 5.2 Hz, 1H), 2.20 – 2.07 (m, 1H).

LC/MS (APSI) m/z [M+H] calculated for C9H13N2: 149.4; found: 149.1.

**2-(naphthalen-1-yl)prop-2-en-1-amine; trifluoroacetic acid – ZINCey000000C7Tw, Z8711777920.**

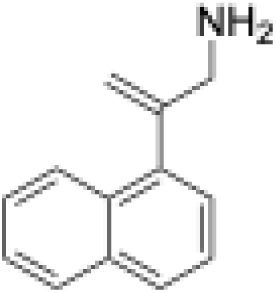

Purity, >95% (assessed by LC/MS).

LC/MS (APSI) m/z [M+H] calculated for C13H14N: 184.2; found: 184.1.

**2-[(pyrrolidin-2-yl)methyl]pyridine dihydrochloride – ZINCcj0000002gSF, Z3214214254.**

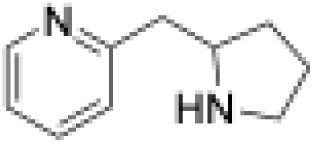

Purity, >95% (assessed by LC/MS).

^1^H NMR (400 MHz, DMSO-*d*_6_) δ 9.68 (s, 2H), 8.81 (d, *J* = 5.6 Hz, 1H), 8.40 (t, *J* = 7.9 Hz, 1H), 8.01 (d, *J* = 7.9 Hz, 1H), 7.83 (t, *J* = 6.7 Hz, 1H), 3.94 (t, *J* = 7.5 Hz, 1H), 3.56 (dd, *J* = 14.6, 8.2 Hz, 1H), 3.47 (dd, *J* = 14.6, 6.2 Hz, 1H), 3.23 (s, 1H), 2.05 (ddd, *J* = 25.3, 12.6, 4.7 Hz, 2H), 1.85 (dt, *J* = 20.7, 7.8 Hz, 1H), 1.76 – 1.62 (m, 1H).

LC/MS (APSI) m/z [M+H] calculated for C10H15N2: 163.2; found: 163.1.

**6-fluoro-3,4-dihydro-2H-1-benzopyran-4-amine hydrochloride – ZINCcm0000002EAW, Z1266854953.**

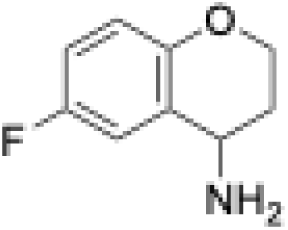

Purity, >95% (assessed by LC/MS).

^1^H NMR (400 MHz, DMSO-*d*_6_) δ 8.76 (s, 3H), 7.47 (dd, *J* = 9.4, 3.2 Hz, 1H), 7.13 (td, *J* = 8.6, 3.1 Hz, 1H), 6.88 (dd, *J* = 9.1, 4.8 Hz, 1H), 4.50 (t, *J* = 5.7 Hz, 1H), 4.23 (dddd, *J* = 17.7, 11.3, 8.8, 3.3 Hz, 2H), 2.24 (tt, *J* = 9.2, 5.0 Hz, 1H), 2.11 (ddt, *J* = 11.7, 6.1, 3.0 Hz, 1H).

LC/MS (APSI) m/z [M+H] calculated for C9H11FNO: 168.0; found: 168.1.

**1-{pyrazolo[1,5-a]pyridin-7-yl}methanamine dihydrochloride – ZINCbd0000002Cxp, Z4244625472.**

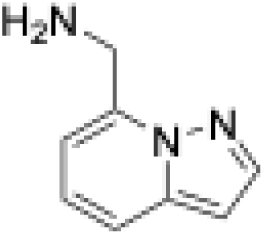

Purity, >95% (assessed by LC/MS).

^1^H NMR (400 MHz, DMSO-*d*_6_) δ 8.83 (s, 3H), 8.12 (d, *J* = 2.2 Hz, 1H), 7.79 (d, *J* = 8.9 Hz, 1H), 7.30 (dd, *J* = 8.8, 6.9 Hz, 1H), 7.12 (d, *J* = 6.9 Hz, 1H), 6.76 (d, *J* = 2.3 Hz, 1H), 4.51 (d, *J* = 5.6 Hz, 2H).

LC/MS (APSI) m/z [M+H] calculated for C8H10N: 148.2; found: 148.1.

**4-[(pyrrolidin-2-yl)methyl]-3,4-dihydro-2H-1,4-benzoxazine dihydrochloride – ZINCgm000000M8UF, Z8700802749.**

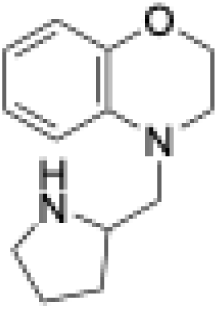

Purity, >95% (assessed by LC/MS).

^1^H NMR (500 MHz, DMSO-*d*_6_) δ 9.35 (s, 1H), 9.14 (s, 1H), 6.81 (dd, *J* = 8.1, 1.5 Hz, 1H), 6.75 (td, *J* = 7.6, 1.6 Hz, 1H), 6.68 (dd, *J* = 7.9, 1.5 Hz, 1H), 6.60 – 6.53 (m, 1H), 4.18 (dtdd, *J* = 10.6, 8.3, 5.5, 3.0 Hz, 2H), 3.71 (t, *J* = 7.5 Hz, 1H), 3.58 (dd, *J* = 14.9, 8.4 Hz, 1H), 3.49 – 3.40 (m, 2H), 3.23 (ddt, *J* = 11.9, 7.6, 4.0 Hz, 1H), 3.10 (ddt, *J* = 15.3, 11.9, 6.2 Hz, 1H), 2.11 (dtd, *J* = 11.8, 7.4, 3.9 Hz, 1H), 1.97 (dq, *J* = 12.8, 4.3 Hz, 1H), 1.89 (tt, *J* = 16.1, 6.1 Hz, 1H), 1.60 (dq, *J* = 12.9, 8.9 Hz, 1H).

LC/MS (APSI) m/z [M+H] calculated for C13H19N2O: 219.2; found: 219.1.

**[(1-benzofuran-3-yl)methyl](methyl)amine – ZINCcr0000000JH8, Z1197877493.**

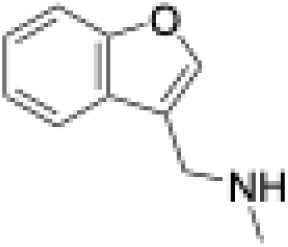

Purity, >95% (assessed by LC/MS).

^1^H NMR (400 MHz, DMSO-*d*_6_) δ 7.82 (s, 1H), 7.71 (dd, *J* = 7.8, 1.4 Hz, 1H), 7.54 (dd, *J* = 8.1, 1.0 Hz, 1H), 7.27 (dtd, *J* = 24.8, 7.2, 1.2 Hz, 2H), 3.76 (s, 2H), 3.30 (s, 1H), 2.31 (s, 3H).

LC/MS (APSI) m/z [M+H] calculated for C10H12NO: 162.0; found: 162.1.

**4-{[(3,4-dihydro-2H-1-benzopyran-4-yl)methyl]amino}but-2-en-1-ol – ZINChm000000Gu6n, Z2590548459.**

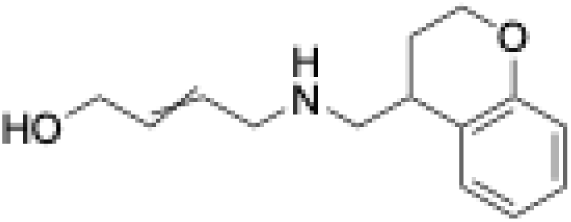

Purity, >95% (assessed by LC/MS).

LC/MS (APSI) m/z [M+H] calculated for C14H20NO2: 234.4; found: 234.1.

**2-(aminomethyl)-3,4-dihydro-2H-1-benzopyran-4-one hydrochloride – ZINCdf000001oqrA, Z2739607300.**

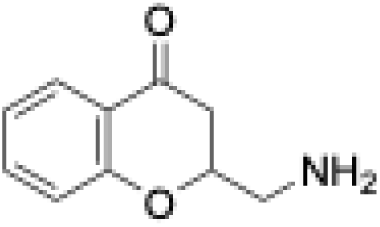

Purity, >95% (assessed by LC/MS).

^1^H NMR (400 MHz, DMSO-*d*_6_) δ 8.47 (s, 3H), 7.78 (d, *J* = 7.8 Hz, 1H), 7.62 (t, *J* = 7.8 Hz, 1H), 7.16 – 7.05 (m, 2H), 4.87 – 4.78 (m, 1H), 3.35 (s, 1H), 2.95 (ddd, *J* = 16.7, 13.1, 3.2 Hz, 1H), 2.77 (dt, *J* = 17.0, 3.3 Hz, 1H).

LC/MS (APSI) m/z [M+H] calculated for C10H12NO2: 178.2; found: 178.1.

**(2R)-1-(7-fluoro-2,3-dihydro-1H-indol-1-yl)propan-2-amine dihydrochloride – ZINCel000000ZhUM, Z8721927756.**

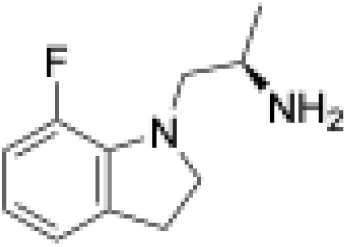

Purity, >95% (assessed by LC/MS).

^1^H NMR (500 MHz, DMSO-*d*_6_) δ 8.03 – 7.99 (m, 3H), 6.93 – 6.83 (m, 2H), 6.63 (td, *J* = 7.7, 4.2 Hz, 1H), 3.45 (q, *J* = 8.7 Hz, 1H), 3.40 – 3.32 (m, 2H), 3.24 (dd, *J* = 13.0, 5.0 Hz, 1H), 2.97 (td, *J* = 8.6, 2.9 Hz, 2H), 1.22 (d, *J* = 6.1 Hz, 3H).

LC/MS (APSI) m/z [M+H] calculated for C11H16FN2: 195.2; found: 195.1.

**[(cyclopent-2-en-1-yl)methyl][(pyridin-2-yl)methyl]amine – ZINCer0000002Srj, Z2787083864.**

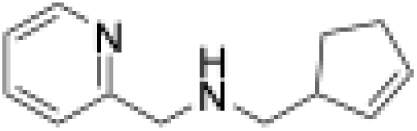

Purity, >95% (assessed by LC/MS).

LC/MS (APSI) m/z [M+H] calculated for C12H17N2: 189.1; found: 189.1.

**1-(2,3-dihydro-1H-inden-1-yl)methanamine hydrochloride – ZINCbm0000002DE2, Z3241259976.**

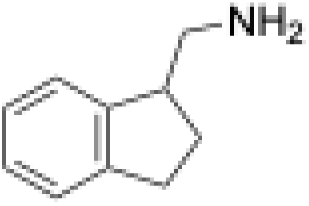

Purity, >95% (assessed by LC/MS).

^1^H NMR (400 MHz, CDCl_3_) δ 7.21 (ddt, *J* = 22.8, 5.6, 3.4 Hz, 4H), 3.22 (p, *J* = 6.7 Hz, 1H), 3.06 – 2.81 (m, 5H), 2.29 (dtd, *J* = 12.6, 8.3, 5.7 Hz, 1H), 1.85 (ddt, *J* = 12.7, 8.7, 6.4 Hz, 1H), 1.15 (s, 2H).

LC/MS (APSI) m/z [M+H] calculated for C10H14N: 148.0; found: 148.1.

**1-(2,3-dihydro-1-benzofuran-3-yl)ethan-1-amine – 7490_27, Z1493816383.**

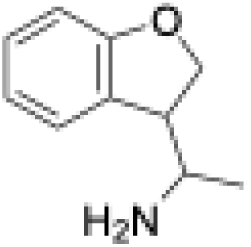

Purity, >95% (assessed by LC/MS).

^1^H NMR (500 MHz, DMSO-*d*_6_) δ 7.28 – 7.22 (m, 1H), 7.07 (t, *J* = 7.6 Hz, 1H), 6.79 (td, *J* = 7.4, 3.7 Hz, 1H), 6.70 (d, *J* = 7.9 Hz, 1H), 4.53 – 4.36 (m, 2H), 3.63 – 2.86 (m, 4H), 0.87 (dd, *J* = 72.6, 6.4 Hz, 3H).

LC/MS (APSI) m/z [M+H] calculated for C10H14NO: 164.1; found: 164.1.

**1-(2,3-dihydro-1-benzothiophen-3-yl)methanamine – 7490_13, Z1316760731.**

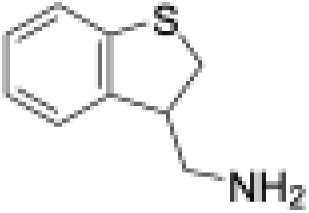

Purity, >95% (assessed by LC/MS).

^1^H NMR (500 MHz, DMSO-*d*_6_) δ 7.19 (dd, *J* = 10.2, 7.6 Hz, 2H), 7.10 (t, *J* = 7.5 Hz, 1H), 7.05 – 6.96 (m, 1H), 3.45 (dd, *J* = 10.6, 7.5 Hz, 1H), 3.41 – 3.27 (m, 2H), 2.78 (dd, *J* = 12.6, 4.8 Hz, 1H), 2.67 (dd, *J* = 12.6, 8.5 Hz, 1H), 1.72 (s, 2H).

LC/MS (APSI) m/z [M+H] calculated for C9H11NS: 166.0; found: 166.1.

**2-(pyridin-3-yl)morpholine dihydrochloride – ZINCcd0000002hfn, Z3235143301.**

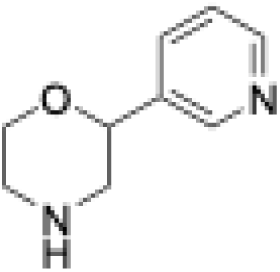

Purity, >95% (assessed by LC/MS).

^1^H NMR (400 MHz, DMSO-*d*_6_) δ 10.11 (s, 2H), 8.87 (dd, *J* = 12.4, 3.8 Hz, 2H), 8.46 (d, *J* = 8.1 Hz, 1H), 7.97 (dd, *J* = 8.1, 5.5 Hz, 1H), 5.10 (dd, *J* = 11.4, 2.6 Hz, 1H), 4.16 (dd, *J* = 12.6, 3.8 Hz, 1H), 4.04 (td, *J* = 12.3, 2.5 Hz, 1H), 3.58 (d, *J* = 12.6 Hz, 1H), 3.25 (d, *J* = 12.8 Hz, 1H), 3.09 (d, *J* = 11.3 Hz, 2H).

LC/MS (APSI) m/z [M+H] calculated for C9H13N2O: 165.2; found: 165.1.

**2-(thiophen-3-yl)morpholine – ZINCbk0000002gVO, Z954552190.**

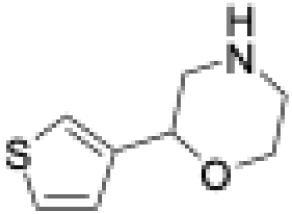

**Purity, >95% (assessed by LC/MS).**

**1H NMR (400 MHz, CDCl3) δ 7.37 – 7.22 (m, 2H), 7.08 (d, J = 5.0 Hz, 1H), 4.60 (dd, J = 10.2, 2.5 Hz, 1H), 4.01 (dd, J = 11.6, 3.3 Hz, 1H), 3.79 (td, J = 11.6, 2.8 Hz, 1H), 3.18 – 3.10 (m, 1H), 3.00 (td, J = 11.9, 3.4 Hz, 1H), 2.98 – 2.83 (m, 2H), 2.25 (s, 1H).**

**LC/MS (APSI) m/z [M+H] calculated for C8H12NOS: 170.2; found: 170.1.**

**2-(pyridin-2-yl)morpholine dihydrochloride – ZINCcd0000002gTu, Z3235143299.**

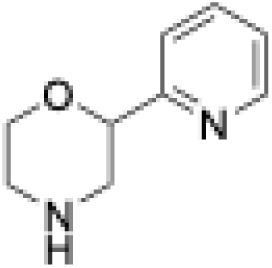

Purity, >95% (assessed by LC/MS).

^1^H NMR (400 MHz, DMSO-*d*_6_) δ 9.99 (s, 1H), 9.76 (s, 1H), 8.63 (d, *J* = 5.1 Hz, 1H), 8.05 (t, *J* = 7.8 Hz, 1H), 7.65 (d, *J* = 7.9 Hz, 1H), 7.54 (dd, *J* = 7.6, 5.0 Hz, 1H), 5.03 (dd, *J* = 11.1, 2.6 Hz, 1H), 4.15 (dd, *J* = 12.5, 3.8 Hz, 1H), 4.03 (td, *J* = 12.3, 2.5 Hz, 1H), 3.60 (d, *J* = 12.6 Hz, 1H), 3.26 (d, *J* = 12.8 Hz, 1H), 3.19 – 2.99 (m, 2H).

LC/MS (APSI) m/z [M+H] calculated for C9H13N2O: 165.0; found: 165.1.

**(4S)-4-(methylamino)-3,4-dihydro-2H-1-benzopyran-6-ol – ZINCdk000002Tr9f, Z3519067312.**

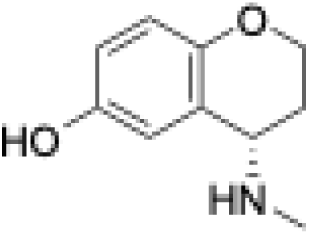

Purity, >95% (assessed by LC/MS).

^1^H NMR (500 MHz, DMSO-*d*_6_) δ 8.71 (s, 1H), 6.67 (d, *J* = 2.1 Hz, 1H), 6.50 (d, *J* = 1.9 Hz, 2H), 4.11 (ddd, *J* = 10.8, 8.2, 4.1 Hz, 1H), 4.03 – 3.96 (m, 1H), 3.47 (t, *J* = 4.8 Hz, 1H), 2.30 (s, 3H), 1.93 (s, 1H), 1.89 – 1.76 (m, 2H).

LC/MS (APSI) m/z [M+H] calculated for C10H14NO2: 180.0; found: 180.1.

**N-methyl-2H,3H,4H-pyrano[3,2-b]pyridin-4-amine – 4953_6, Z1262608488.**

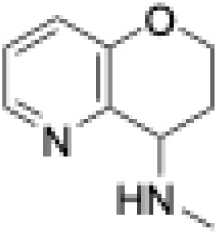

Purity, >95% (assessed by LC/MS).

LC/MS (APSI) m/z [M+H] calculated for C9H13N2O: 165.0; found: 165.1.

**3,4-dihydro-2H-1-benzopyran-4-amine – 4953_4, Z85923435.**

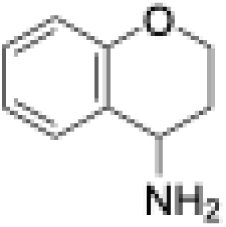

Purity, >95% (assessed by LC/MS).

^1^H NMR (400 MHz, CDCl_3_) δ 7.37 – 7.10 (m, 2H), 6.92 (td, *J* = 7.4, 1.2 Hz, 1H), 6.83 (dd, *J* = 8.2, 1.2 Hz, 1H), 4.36 – 4.19 (m, 2H), 4.07 (t, *J* = 5.3 Hz, 1H), 2.24 – 2.12 (m, 1H), 1.87 (dtd, *J* = 14.2, 5.8, 2.9 Hz, 1H), 1.71 (d, *J* = 3.7 Hz, 2H).

LC/MS (APSI) m/z [M] calculated for C9H11NO: 149.0; found: 149.1.

**N-methyl-5,6,7,8-tetrahydroquinolin-8-amine – ZINCcm0000002wLB, Z1262565741.**

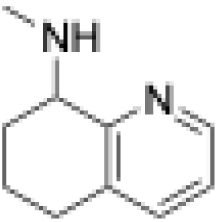

Purity, >95% (assessed by LC/MS).

^1^H NMR (400 MHz, CDCl_3_) δ 8.37 (d, *J* = 4.8 Hz, 1H), 7.35 (d, *J* = 7.7 Hz, 1H), 7.04 (dd, *J* = 7.7, 4.7 Hz, 1H), 3.64 (t, *J* = 6.2 Hz, 1H), 2.86 – 2.67 (m, 2H), 2.59 (s, 1H), 2.52 (s, 3H), 2.12 (d, *J* = 15.8 Hz, 1H), 1.97 (dt, *J* = 12.7, 5.6 Hz, 1H), 1.82 – 1.67 (m, 2H).

LC/MS (APSI) m/z [M+H] calculated for C10H15N2: 163.2; found: 163.1.

**3-[(pyrrolidin-2-yl)methyl]pyridine – ZINCcj0000002hf5, Z1198160843.**

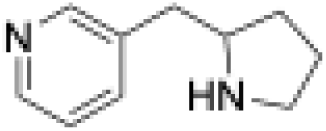

Purity, >95% (assessed by LC/MS).

^1^H NMR (400 MHz, CDCl_3_) δ 8.46 – 8.39 (m, 2H), 7.51 (dt, *J* = 7.8, 2.1 Hz, 1H), 7.17 (dd, *J* = 7.8, 4.8 Hz, 1H), 3.19 (p, *J* = 7.0 Hz, 1H), 2.99 (ddd, *J* = 10.3, 7.6, 5.1 Hz, 1H), 2.80 (ddd, *J* = 10.4, 8.2, 6.6 Hz, 1H), 2.70 (d, *J* = 6.9 Hz, 2H), 1.88 – 1.61 (m, 4H), 1.41 – 1.28 (m, 1H).

LC/MS (APSI) m/z [M+H] calculated for C10H15N2: 163.4; found: 163.1.

**2-[(3-fluorophenyl)methyl]pyrrolidine hydrochloride – 4732_2, Z449369104.**

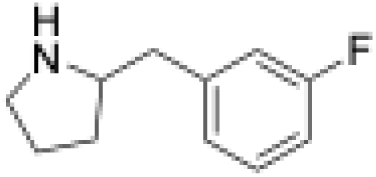

Purity, >95% (assessed by LC/MS).

^1^H NMR (400 MHz, DMSO-*d*_6_) δ 9.27 (s, 2H), 7.38 (td, *J* = 7.9, 6.2 Hz, 1H), 7.26 – 7.14 (m, 2H), 7.09 (td, *J* = 8.7, 2.7 Hz, 1H), 3.66 (p, *J* = 7.0 Hz, 1H), 3.34 (s, 1H), 3.21 (s, 1H), 3.12 (dd, *J* = 13.9, 7.4 Hz, 1H), 2.96 (dd, *J* = 13.8, 7.3 Hz, 1H), 2.04 – 1.76 (m, 2H), 1.60 (dq, *J* = 12.5, 8.2 Hz, 1H).

LC/MS (APSI) m/z [M+H] calculated for C11H15FN: 180.2; found: 180.1.

**(2S)-2-benzylpyrrolidine hydrochloride – ZINCcp0000002gSD, Z3244742843.**

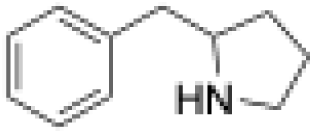

Purity, >95% (assessed by LC/MS).

^1^H NMR (400 MHz, DMSO-*d*_6_) δ 9.34 (d, *J* = 30.0 Hz, 2H), 7.32 (s, 3H), 7.38 – 7.21 (m, 2H), 3.63 (s, 1H), 3.35 (s, 1H), 3.11 (dt, *J* = 18.7, 9.3 Hz, 2H), 2.91 (dd, *J* = 13.6, 7.9 Hz, 1H), 1.95 (s, 1H), 1.96 – 1.81 (m, 1H), 1.84 (s, 1H), 1.60 (dq, *J* = 12.0, 8.9 Hz, 1H).

LC/MS (APSI) m/z [M+H] calculated for C11H16N: 162.2; found: 162.1.

**methyl[(quinolin-8-yl)methyl]amine hydrochloride – ZINCdp000000Y2uG, Z4710975379.**

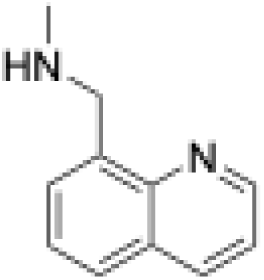

Purity, >95% (assessed by LC/MS).

^1^H NMR (400 MHz, DMSO-*d*_6_) δ 9.48 (s, 2H), 9.06 (dd, *J* = 4.5, 1.7 Hz, 1H), 8.63 (dq, *J* = 6.6, 2.1 Hz, 1H), 8.15 (dd, *J* = 8.3, 1.4 Hz, 1H), 8.07 (d, *J* = 7.1 Hz, 1H), 7.79 – 7.69 (m, 2H), 4.74 (t, *J* = 5.9 Hz, 2H), 2.61 (td, *J* = 5.6, 2.0 Hz, 3H).

LC/MS (APSI) m/z [M+H] calculated for C11H13N2: 173.2; found: 173.1.

**1-(quinolin-2-yl)methanamine dihydrochloride – ZINCcm0000002CuJ, Z2106596235.**

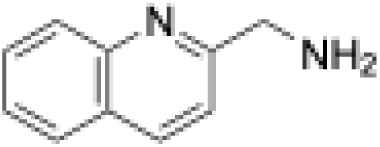

Purity, >95% (assessed by LC/MS).

^1^H NMR (400 MHz, DMSO-*d*_6_) δ 8.80 (s, 3H), 8.56 (d, *J* = 8.5 Hz, 1H), 8.10 (dd, *J* = 14.7, 8.3 Hz, 2H), 7.88 (ddd, *J* = 8.5, 6.8, 1.4 Hz, 1H), 7.76 (d, *J* = 8.5 Hz, 1H), 7.70 (t, *J* = 7.5 Hz, 1H), 4.44 (q, *J* = 5.8 Hz, 2H).

LC/MS (APSI) m/z [M+H] calculated for C10H11N2: 159.3; found: 159.1.

**[2-(isoquinolin-1-yl)prop-2-en-1-yl](methyl)amine triflate – 7920_1, Z8843458836.**

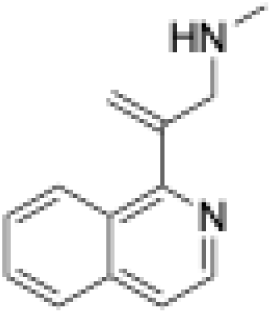

Purity, >95% (assessed by LC/MS).

LC/MS (APSI) m/z [M+H] calculated for C13H15N2: 199.0; found: 199.1.

**1-(quinolin-8-yl)ethan-1-amine – ZINCds000000IYAl, Z1198724030.**

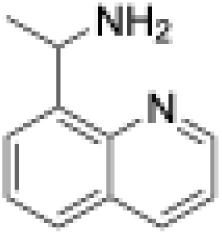

Purity, >95% (assessed by LC/MS).

^1^H NMR (400 MHz, CDCl_3_) δ 8.92 (dd, *J* = 4.2, 1.9 Hz, 1H), 8.15 (dd, *J* = 8.2, 1.9 Hz, 1H), 7.78 – 7.67 (m, 2H), 7.52 (t, *J* = 7.6 Hz, 1H), 7.40 (dd, *J* = 8.3, 4.2 Hz, 1H), 5.18 (q, *J* = 6.7 Hz, 1H), 1.97 (s, 3H), 1.61 (d, *J* = 6.6 Hz, 3H).

LC/MS (APSI) m/z [M+H] calculated for C11H13N2: 173.2; found: 173.1.

**1-(quinolin-8-yl)methanamine – ZINCcm0000002Cxl, Z234897549.**

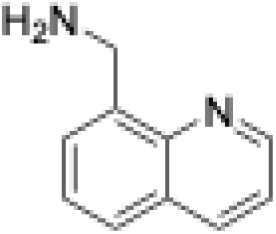

Purity, >95% (assessed by LC/MS).

^1^H NMR (400 MHz, CDCl_3_) δ 8.91 (dd, *J* = 4.2, 1.8 Hz, 1H), 8.14 (dd, *J* = 8.3, 1.8 Hz, 1H), 7.70 (dd, *J* = 8.2, 1.5 Hz, 1H), 7.62 (d, *J* = 6.9 Hz, 1H), 7.47 (dd, *J* = 8.2, 7.0 Hz, 1H), 7.39 (dd, *J* = 8.3, 4.2 Hz, 1H), 4.40 (s, 2H), 1.92 (d, *J* = 8.4 Hz, 2H).

LC/MS (APSI) m/z [M+H] calculated for C10H11N2: 159.2; found: 159.1.

**1-(isoquinolin-1-yl)methanamine dihydrochloride – ZINCcm0000002D5q, Z220439492.**

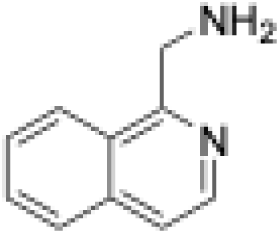

Purity, >95% (assessed by LC/MS).

^1^H NMR (400 MHz, DMSO-*d*_6_) δ 8.72 (s, 3H), 8.54 (d, *J* = 5.8 Hz, 1H), 8.29 (d, *J* = 8.4 Hz, 1H), 8.09 (d, *J* = 8.2 Hz, 1H), 7.95 (d, *J* = 5.8 Hz, 1H), 7.90 (t, *J* = 7.5 Hz, 1H), 7.79 (t, *J* = 7.6 Hz, 1H), 4.80 (q, *J* = 5.8 Hz, 2H).

LC/MS (APSI) m/z [M+H] calculated for C10H11N2: 159.2; found: 159.1.

**2-(pyridin-2-yl)morpholine dihydrochloride – Z8987163207, Z8987163207.**

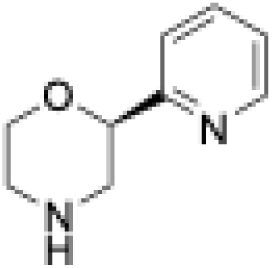

Purity, >95% (assessed by LC/MS).

^1^H NMR (400 MHz, DMSO-*d*_6_) δ 10.45 – 9.51 (m, 2H), 8.63 (d, *J* = 5.2 Hz, 1H), 8.09 – 8.01 (m, 1H), 7.65 (d, *J* = 8.0 Hz, 1H), 7.54 (dd, *J* = 7.6, 5.1 Hz, 1H), 5.03 (dd, *J* = 11.1, 2.5 Hz, 1H), 4.15 (dd, *J* = 12.5, 3.8 Hz, 1H), 4.03 (td, *J* = 12.3, 2.6 Hz, 1H), 3.60 (d, *J* = 12.6 Hz, 1H), 3.25 (s, 1H), 3.11 (dd, *J* = 16.1, 6.9 Hz, 2H).

LC/MS (APSI) m/z [M+H] calculated for C9H13N2O: 165.0; found: 165.1.

**3-[(methylamino)methyl]-2,3-dihydro-1H-inden-4-ol hydrochloride – 7490_22, Z2732542387.**

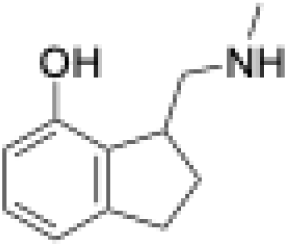

Purity, >95% (assessed by LC/MS).

^1^H NMR (500 MHz, DMSO-*d*_6_) δ 9.70 (s, 1H), 8.79 (s, 2H), 6.99 (t, *J* = 7.7 Hz, 1H), 6.65 (dd, *J* = 13.7, 7.7 Hz, 2H), 3.52 (tt, *J* = 8.9, 4.4 Hz, 1H), 3.27 – 3.21 (m, 1H), 2.90 (dt, *J* = 15.9, 7.9 Hz, 1H), 2.84 (d, *J* = 11.3 Hz, 1H), 2.75 (ddd, *J* = 16.0, 8.8, 4.7 Hz, 1H), 2.56 (s, 3H), 2.12 (dq, *J* = 13.1, 8.1 Hz, 1H), 2.02 (ddt, *J* = 13.0, 8.7, 4.5 Hz, 1H).

LC/MS (APSI) m/z [M+H] calculated for C11H16NO: 178.2; found: 178.1.

**[(3,4-dihydro-1H-2-benzopyran-1-yl)methyl](methyl)amine hydrochloride – ZINCdl000001of0A, Z337709402.**

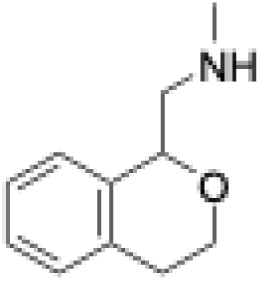

Purity, >95% (assessed by LC/MS).

^1^H NMR (400 MHz, DMSO-*d*_6_) δ 9.32 (s, 1H), 8.77 (s, 1H), 7.21 (q, *J* = 5.5 Hz, 4H), 5.14 – 5.07 (m, 1H), 4.07 (dt, *J* = 10.7, 5.0 Hz, 1H), 3.78 (ddd, *J* = 11.8, 7.8, 4.3 Hz, 1H), 3.48 (d, *J* = 13.0 Hz, 1H), 3.19 (t, *J* = 11.6 Hz, 1H), 2.92 – 2.80 (m, 1H), 2.75 (dt, *J* = 16.5, 4.8 Hz, 1H), 2.58 (s, 3H).

LC/MS (APSI) m/z [M+H] calculated for C11H16NO: 178.1; found: 178.1.

**1-(3,4-dihydro-1H-2-benzopyran-1-yl)methanamine hydrochloride – ZINCci0000002DUp, Z337709400.**

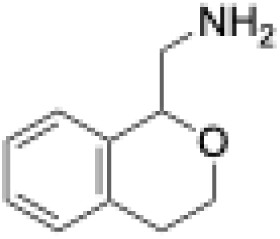

Purity, >95% (assessed by LC/MS).

^1^H NMR (400 MHz, DMSO-*d*_6_) δ 8.19 (s, 3H), 7.28 – 7.15 (m, 4H), 4.98 (dd, *J* = 10.0, 2.8 Hz, 1H), 4.08 (dt, *J* = 11.3, 5.0 Hz, 1H), 3.77 (ddd, *J* = 11.8, 8.1, 4.3 Hz, 1H), 3.42 – 3.30 (m, 1H), 3.05 (dd, *J* = 13.2, 9.8 Hz, 1H), 2.87 (ddd, *J* = 16.6, 8.1, 5.1 Hz, 1H), 2.74 (dt, *J* = 16.5, 4.7 Hz, 1H).

LC/MS (APSI) m/z [M+H] calculated for C10H14NO: 164.0; found: 164.1.

**2-(1,2,3,4-tetrahydronaphthalen-1-yl)pyrrolidine hydrochloride – ZINCfy000000Jliy, Z995211376.**

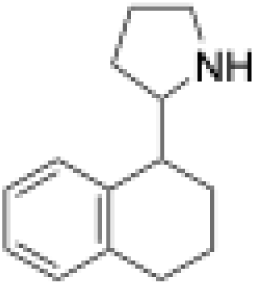

Purity, >95% (assessed by LC/MS).

^1^H NMR (500 MHz, DMSO-*d*_6_) δ 9.48 (s, 1H), 8.98 (s, 1H), 7.39 – 6.88 (m, 4H), 3.58 (s, 1H), 3.20 – 3.08 (m, 3H), 2.85 – 2.76 (m, 1H), 2.75 – 2.65 (m, 1H), 2.06 (d, *J* = 1.7 Hz, 2H), 1.98 – 1.89 (m, 2H), 1.88 – 1.71 (m, 2H), 1.72 (s, 2H).

LC/MS (APSI) m/z [M+H] calculated for C14H20N: 202.2; found: 202.2.

**1-(2,3-dihydro-1-benzothiophen-3-yl)methanamine hydrochloride – EN300-47307925, EN300-47307925.**

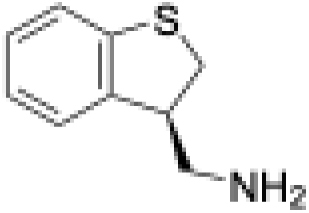

Purity, >95% (assessed by LC/MS).

^1^H NMR (500 MHz, DMSO-*d*_6_) δ 8.28 (s, 3H), 7.30 (d, *J* = 7.5 Hz, 1H), 7.25 (d, *J* = 7.7 Hz, 1H), 7.18 (td, *J* = 7.5, 1.2 Hz, 1H), 7.07 (td, *J* = 7.5, 1.2 Hz, 1H), 3.78 (dq, *J* = 14.4, 5.2 Hz, 1H), 3.54 (dd, *J* = 11.6, 7.5 Hz, 1H), 3.37 (dd, *J* = 11.5, 5.5 Hz, 1H), 3.10 (dd, *J* = 12.8, 4.3 Hz, 1H), 2.96 (dd, *J* = 12.8, 10.0 Hz, 1H).

LC/MS (APSI) m/z [M+H] calculated for C9H12NS: 166.0; found: 166.1.

**(2R)-2-phenylmorpholine hydrochloride – Z1255382255, Z1255382255.**

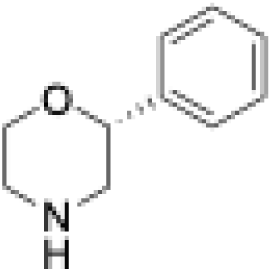

Purity, >95% (assessed by LC/MS).

^1^H NMR (400 MHz, DMSO-*d*_6_) δ 9.72 (s, 2H), 7.38 (s, 3H), 7.44 – 7.31 (m, 2H), 4.80 (dt, *J* = 11.3, 2.2 Hz, 1H), 4.14 – 4.06 (m, 1H), 4.02 – 3.90 (m, 1H), 3.38 (d, *J* = 12.6 Hz, 1H), 3.22 (s, 1H), 3.09 (tt, *J* = 11.9, 3.3 Hz, 1H), 3.02 – 2.91 (m, 1H).

LC/MS (APSI) m/z [M+H] calculated for C10H14NO: 164.0; found: 164.1.

**rel-2-[(1R)-1,2,3,4-tetrahydronaphthalen-1-yl]pyrrolidine hydrochloride – Z8987197482, Z8987197482.**

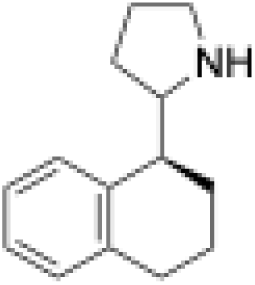

Purity, >95% (assessed by LC/MS).

^1^H NMR (500 MHz, DMSO-*d*_6_) δ 9.33 (s, 2H), 7.16 (dd, *J* = 23.9, 7.4 Hz, 2H), 7.09 (dd, *J* = 10.2, 7.5 Hz, 2H), 3.57 (td, *J* = 9.8, 5.0 Hz, 1H), 3.15 (qd, *J* = 5.6, 3.4 Hz, 2H), 3.11 (dd, *J* = 9.6, 4.8 Hz, 1H), 2.80 (ddd, *J* = 17.0, 6.8, 3.7 Hz, 1H), 2.69 (dt, *J* = 16.7, 8.2 Hz, 1H), 2.01 – 1.64 (m, 7H).

LC/MS (APSI) m/z [M+H] calculated for C14H20N: 202.2; found: 202.2.

**2-phenylmorpholine – ZINCcj0000002gTq, Z283821656.**

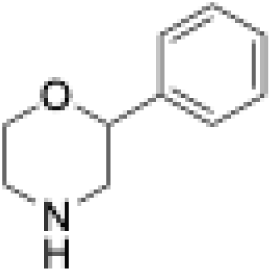

Purity, >95% (assessed by LC/MS).

^1^H NMR (400 MHz, CDCl_3_) δ 7.41 – 7.16 (m, 5H), 4.47 (dd, *J* = 10.3, 2.6 Hz, 1H), 4.02 (dd, *J* = 11.5, 3.4 Hz, 1H), 3.76 (td, *J* = 11.5, 2.7 Hz, 1H), 3.08 – 2.92 (m, 2H), 2.91 – 2.84 (m, 1H), 2.78 (dd, *J* = 12.4, 10.2 Hz, 1H), 1.93 (s, 1H).

LC/MS (APSI) m/z [M+H] calculated for C10H14NO: 164.0; found: 164.1.

## Extended Data

**Extended Data Fig. 1.**
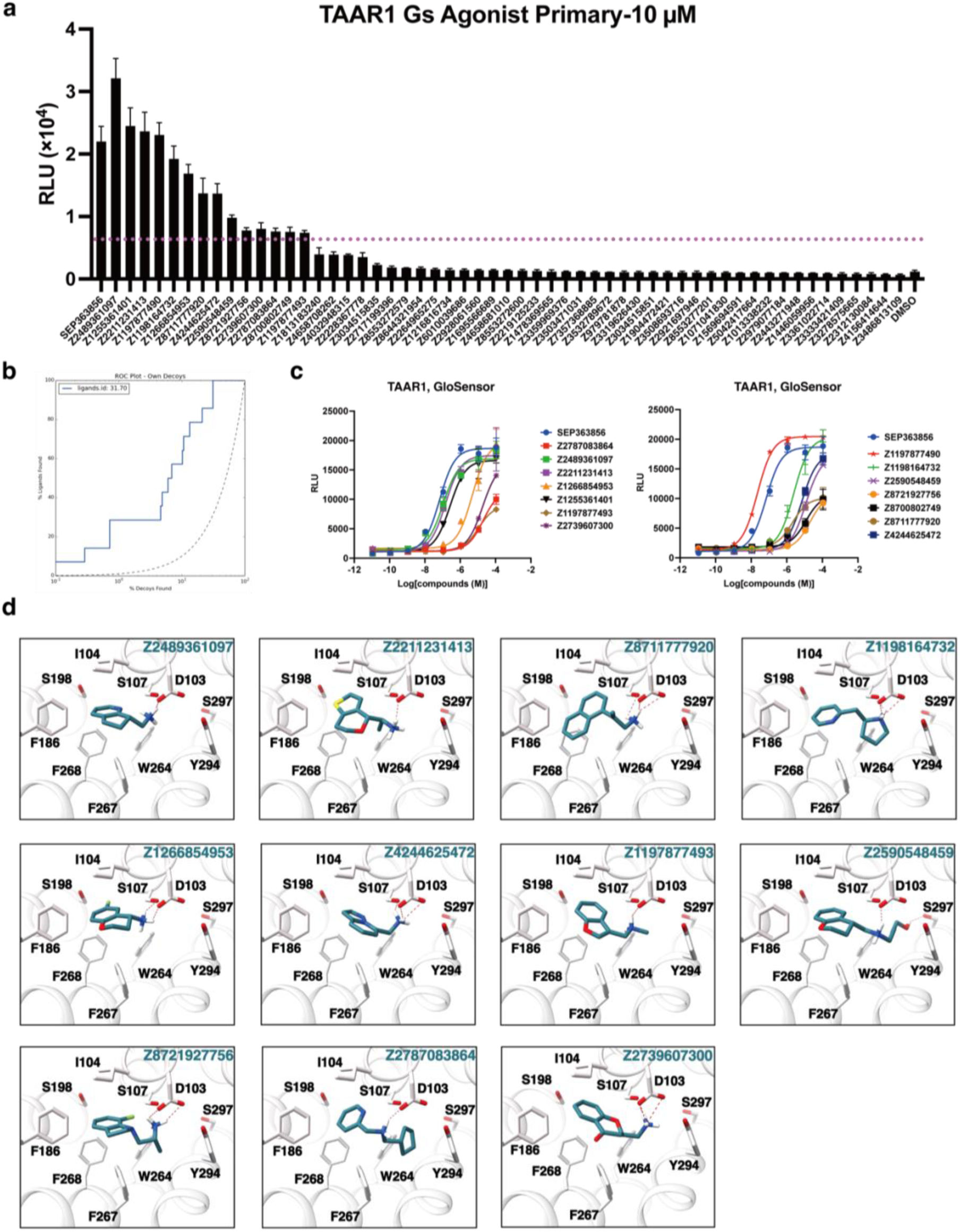
Functional screening of the predicted molecules against the TAAR1 and the docked poses of the initial hits from the larger screen. **a,** GloSensor responses of 55 predicted molecules at 10 μM. Hit rates were defined as having more than a 30% GloSensor response compared with SEP-363856 at 10 μM. **b,** Log-transformed ROC plots that compare the rate of ligand identification versus property-matched decoys for the TAAR1 receptor. **c,** GloSensor response of the initial hits at 10 μM. **d,** Docked poses of the initial hits from the 65-million screen of compounds.

**Extended Data Fig. 2.**
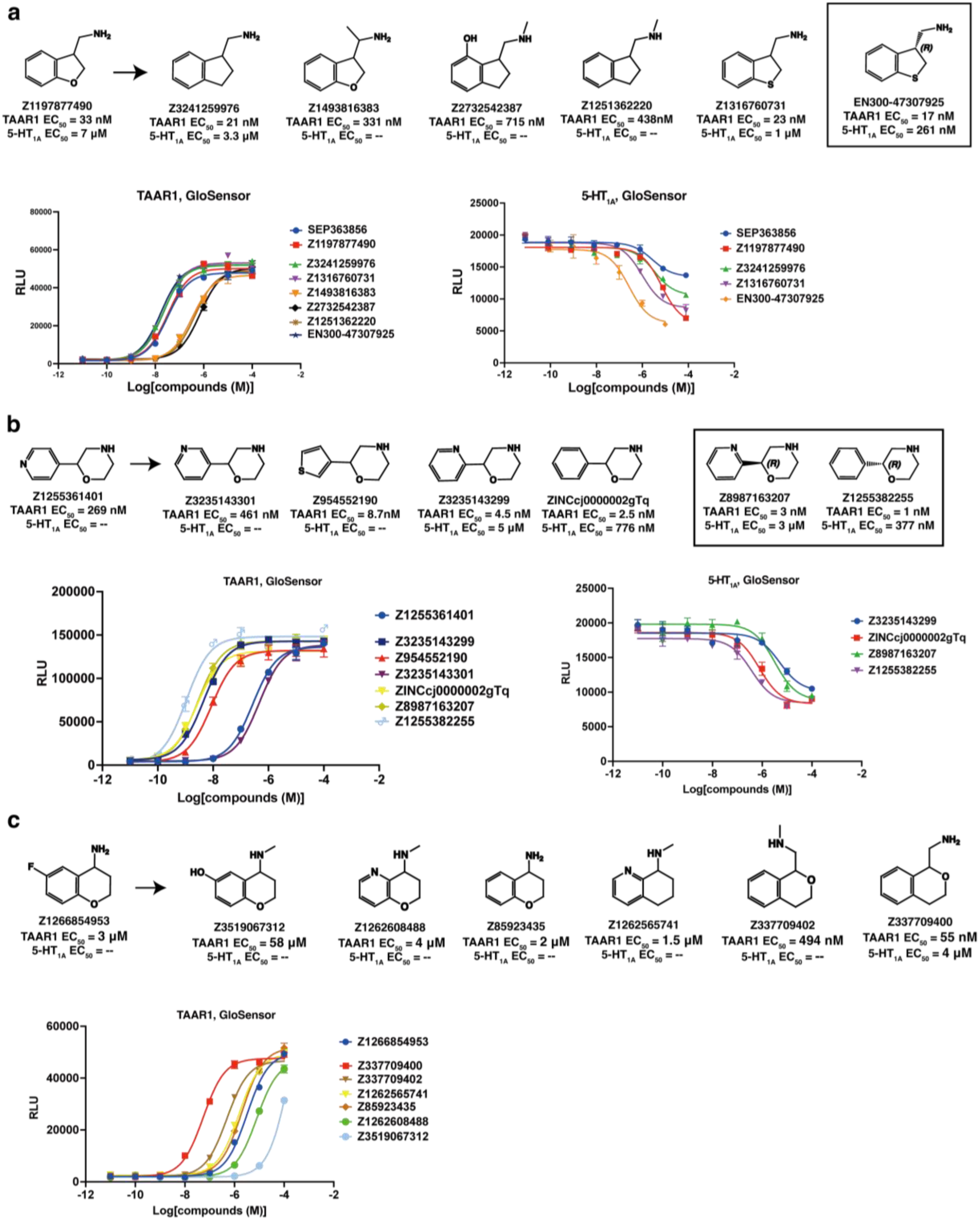
Additional structure-activity relationships around the initial hits from a large-scale docking campaign. **a,** Additional SAR for optimization of compound ‘7490. **b,** Additional SAR for optimization of compound ‘1401. **c,** SAR for optimization of compound ‘4953.

**Extended Data Fig. 3.**
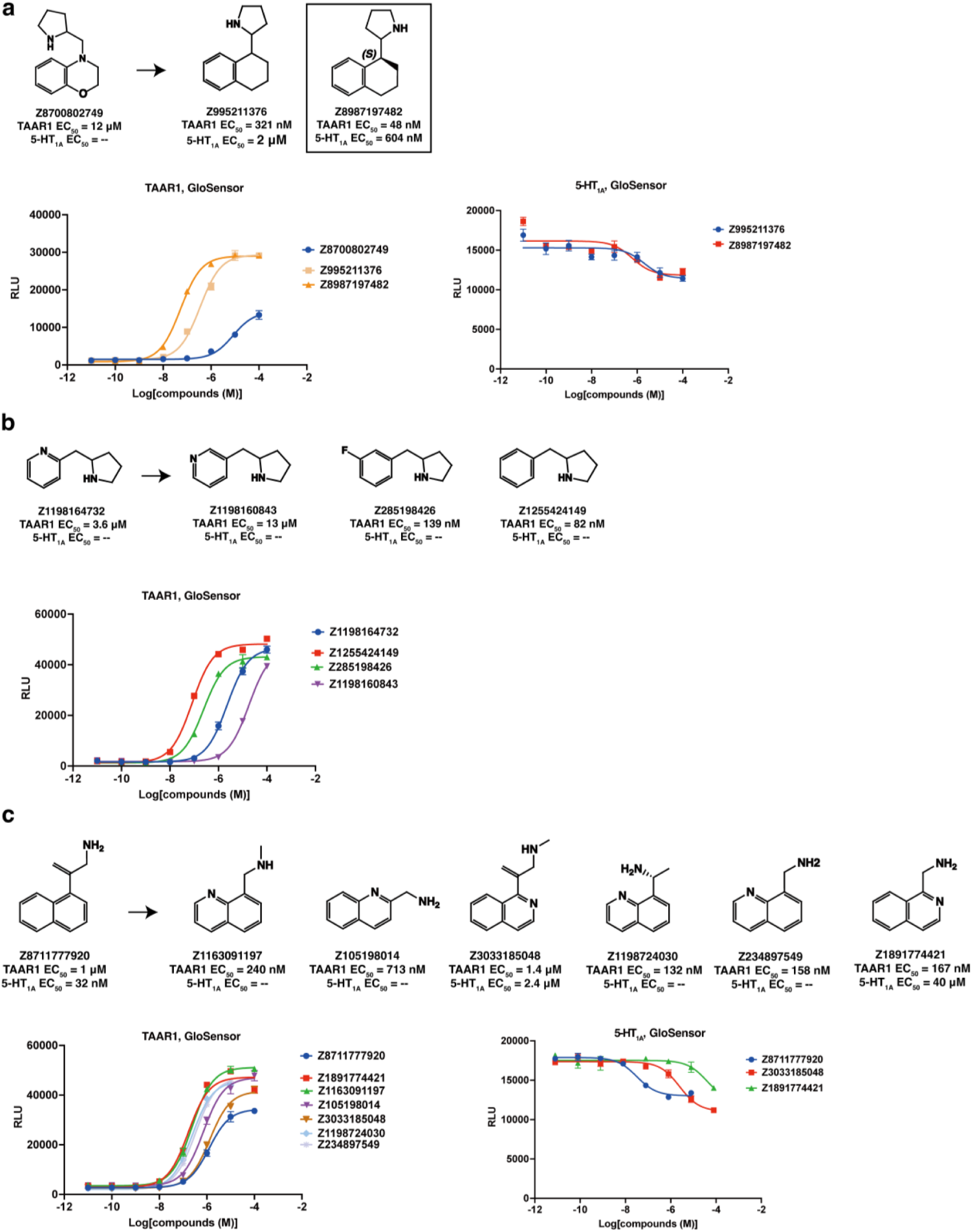
Additional structure-activity relationships around the initial hits from a large-scale docking campaign. **a,** Additional SAR for optimization of compound ‘2749. **b,** SAR for optimization of compound ‘4732. **c,** SAR for optimization of compound ‘7920.

**Extended Data Fig. 4.**
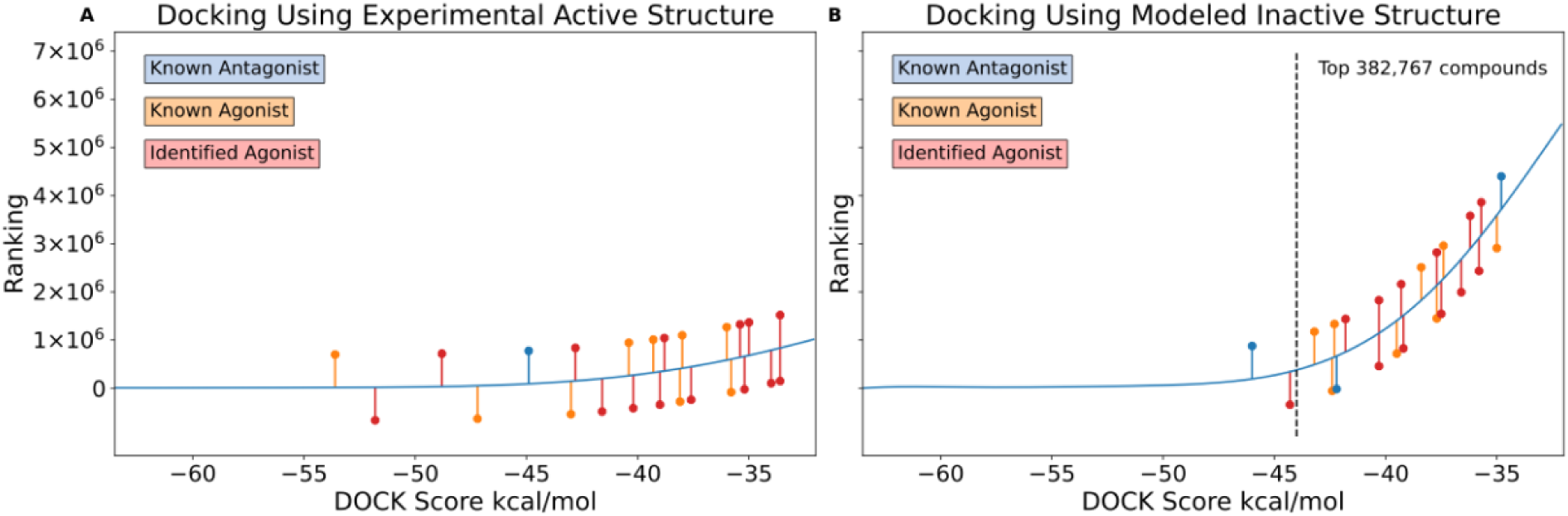
Comparison of large-scale docking results using the experimentally active and modeled inactive TAAR1 structures. **a,** Docking results using the experimental active structure, showing the rankings and docking scores of known agonists, antagonists and identified agonists. **b,** Docking results using the modeled inactive structure (generated by Boltz-2), showing the rankings and docking scores of agonists and antagonists. The known antagonist, known agonist and identified agonist are shown in blue, orange and red respectively. The top 382,767 ranking position is labelled (b).

**Extended Data Fig. 5.**
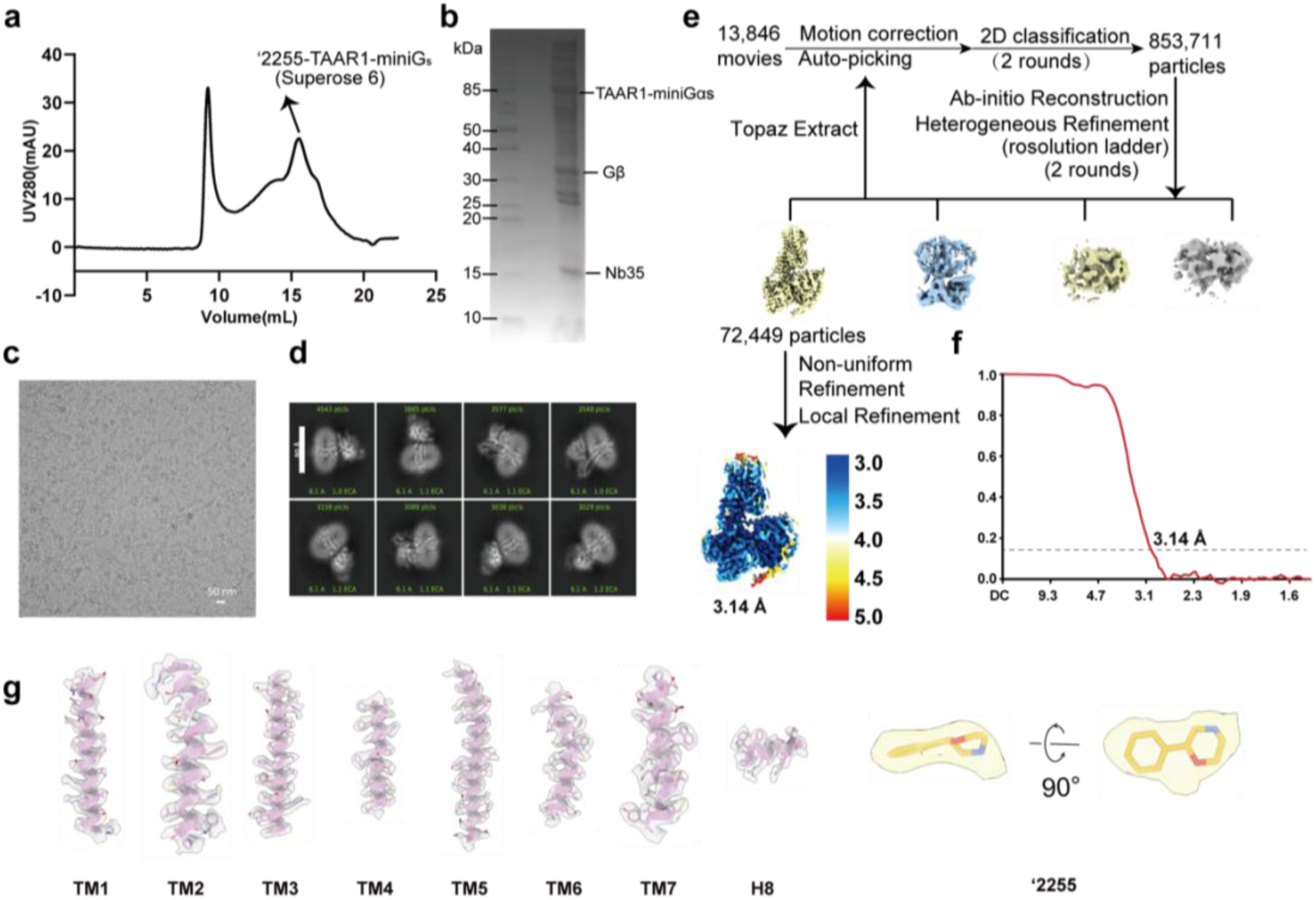
Purification and structural determination of ‘2255 bound TAAR1-Gs complexes. **a,** Representative size-exclusion chromatography (SEC) profiles. **b,** SDS-PAGE analysis of the TAAR1-Gs complex activated by ‘2255. **c-d,** Representative cryo-EM image from 13,846 movies **(c)** and 2D classification averages **(d)** of ‘2255-TAAR1-Gs. **e,** Cryo-EM data-processing flowcharts of ‘2255-TAAR1-Gs by cyroSPARC 3.2 and the global density map of ‘2255-TAAR1-Gs colored by local resolutions. **f,** The Fourier shell correlation (FSC) curves of ‘2255-TAAR1-Gs. Global resolution of the final processed density map estimated at the FSC = 0.143 is 3.14 Å. **g,** The density maps of helices TM1-TM7 of the transmembrane domain and extracellular loop ECL2 of TAAR1, as well as the α5 helix of Gαs with ‘2255.

**Extended Data Fig. 6.**
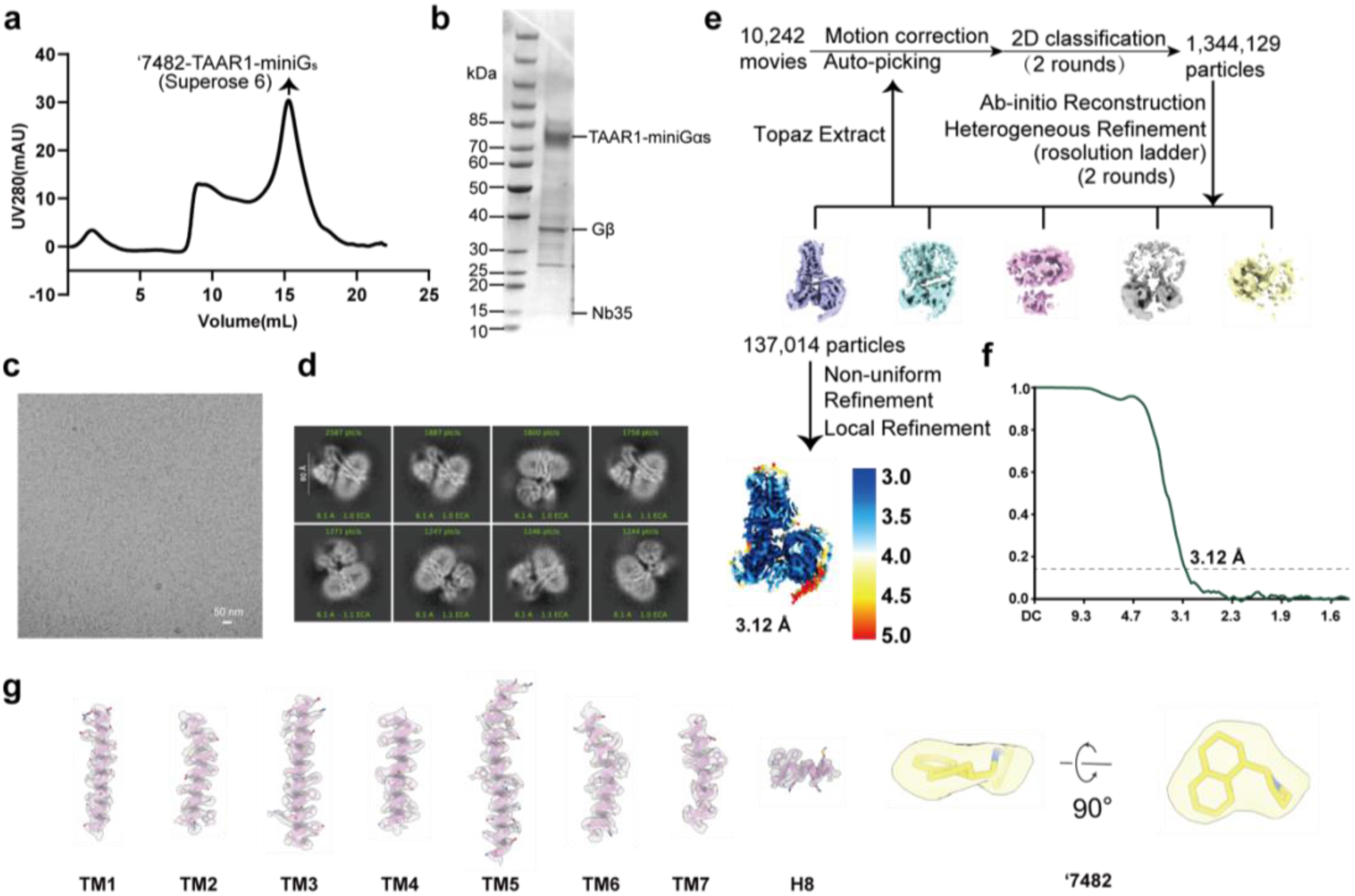
Purification and structural determination of ‘7482 bound TAAR1-Gs complexes. **a,** Representative size-exclusion chromatography (SEC) profiles. **b,** SDS-PAGE analysis of TAAR1-Gs complex activated by ‘7482. **c-d,** Representative cryo-EM image from 10,242 movies **(c)** and 2D classification averages **(d)** of ‘7482-TAAR1-Gs. **e,** Cryo-EM data-processing flowcharts of ‘7482-TAAR1-Gs by cyroSPARC 3.2 and the global density map of ‘7482-TAAR1-Gs colored by local resolutions. **f,** The Fourier shell correlation (FSC) curves for ‘7482-TAAR1-Gs. Global resolution of the final processed density map estimated at the FSC = 0.143 is 3.12 Å. **g,** The density maps of helices TM1-TM7 of the transmembrane domain and extracellular loop ECL2 of TAAR1, as well as the α5 helix of Gαs with ‘7482.

**Extended Data Fig. 7.**
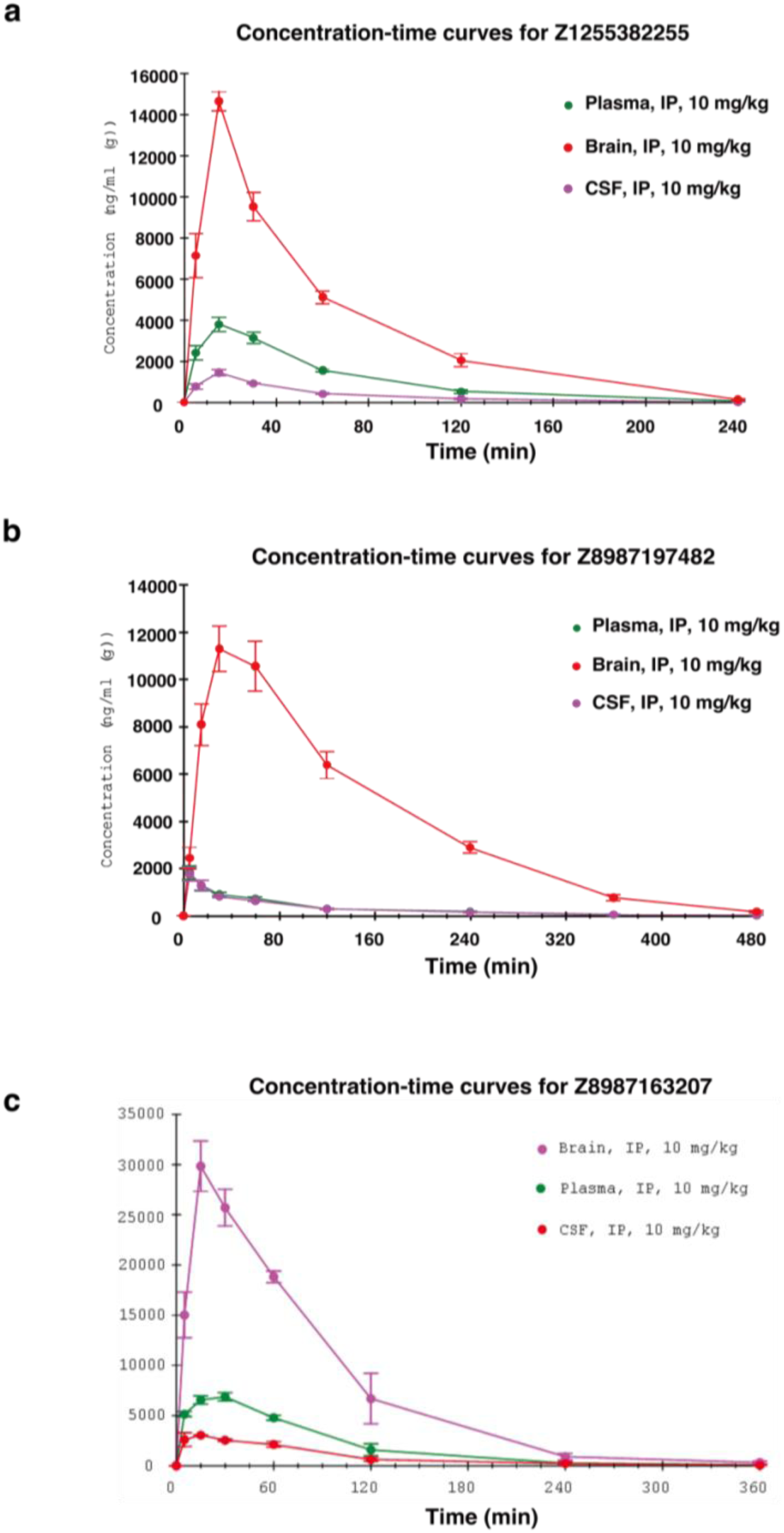
Pharmacokinetic profile of compounds in mice. **a,** Concentration-time curves of **‘2255** in the blood plasma, brain, and cerebrospinal fluid (CSF) after i.p. administration at 10 mg/kg to mice. **b,** Concentration-time curves of **‘7482** in the blood plasma, brain, and cerebrospinal fluid (CSF) after i.p. administration at 10 mg/kg to mice. **c,** Concentration-time curves of **‘3207** in the blood plasma, brain, and cerebrospinal fluid (CSF) after i.p. administration at 10 mg/kg to mice.

**Extended Data Table 1.**
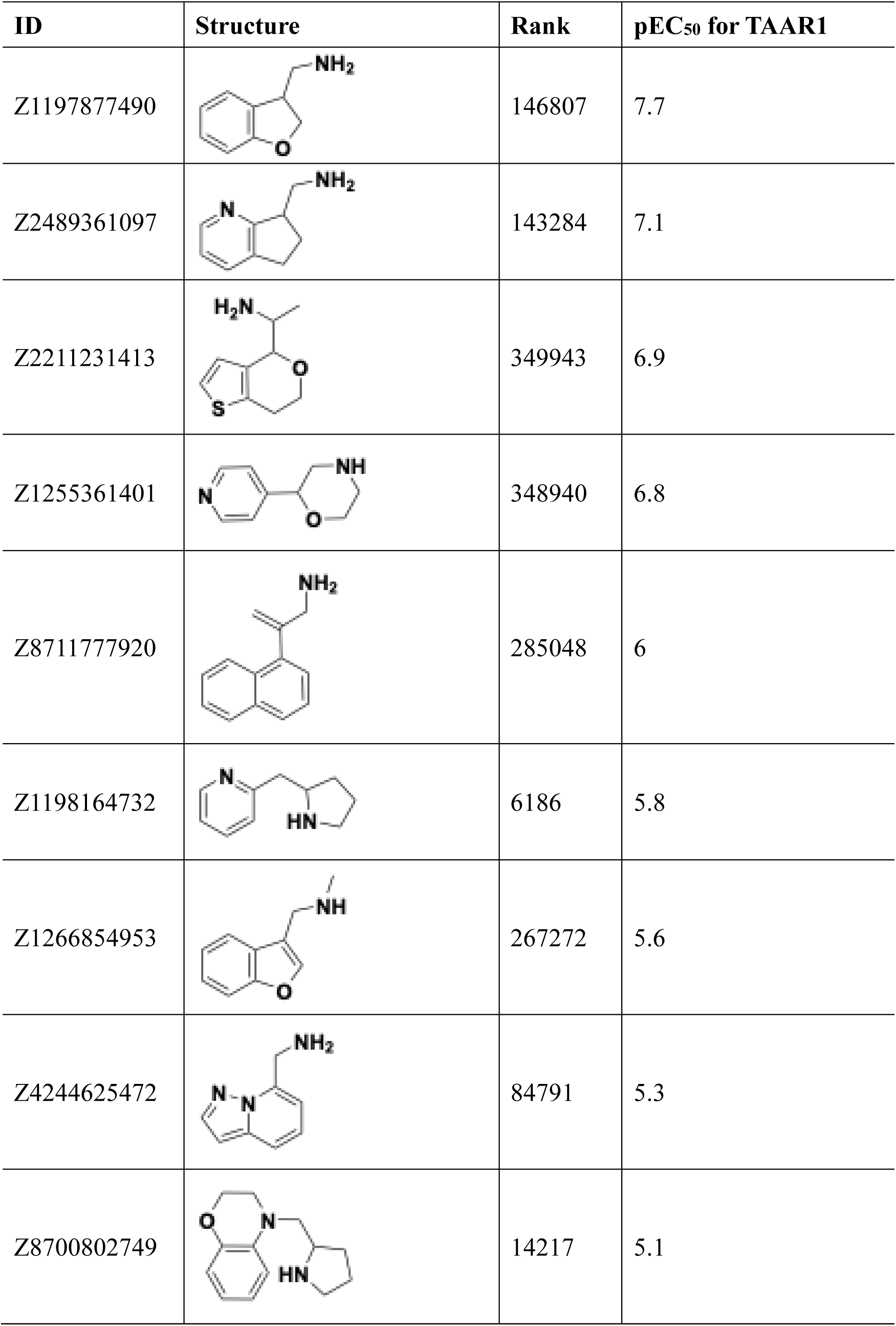
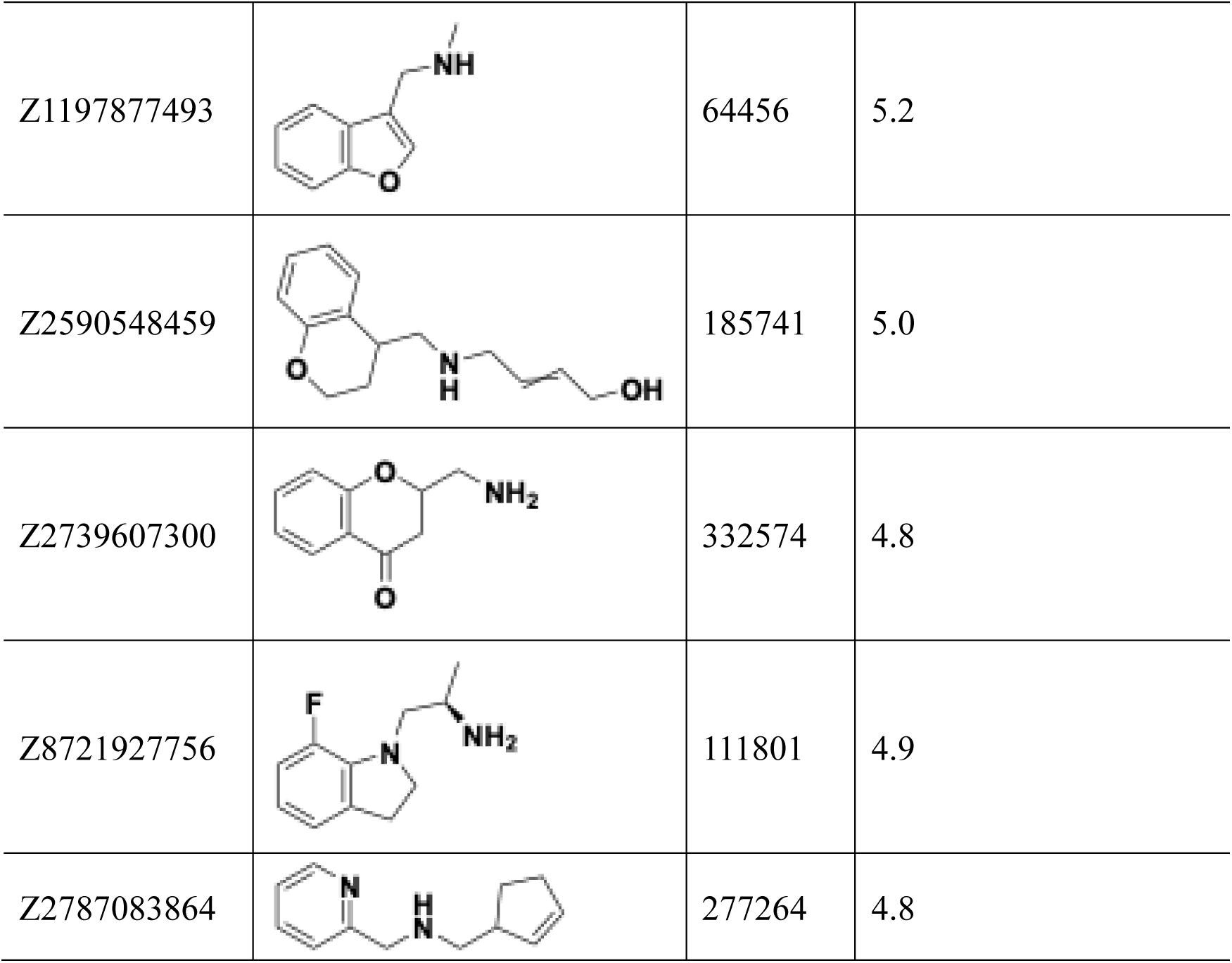
Discovered initial hits from the docking screen against TAAR1.

| ID | Structure | Rank | pEC<sub>50</sub> for TAAR1 |
| --- | --- | --- | --- |
| Z1197877490 |  | 146807 | 7.7 |
| Z2489361097 |  | 143284 | 7.1 |
| Z2211231413 |  | 349943 | 6.9 |
| Z1255361401 |  | 348940 | 6.8 |
| Z8711777920 |  | 285048 | 6 |
| Z1198164732 |  | 6186 | 5.8 |
| Z1266854953 |  | 267272 | 5.6 |
| Z4244625472 |  | 84791 | 5.3 |
| Z8700802749 |  | 14217 | 5.1 |
| Z1197877493 |  | 64456 | 5.2 |
| Z2590548459 |  | 185741 | 5.0 |
| Z2739607300 |  | 332574 | 4.8 |
| Z8721927756 |  | 111801 | 4.9 |
| Z2787083864 |  | 277264 | 4.8 |

**Extended Data Table 2.**
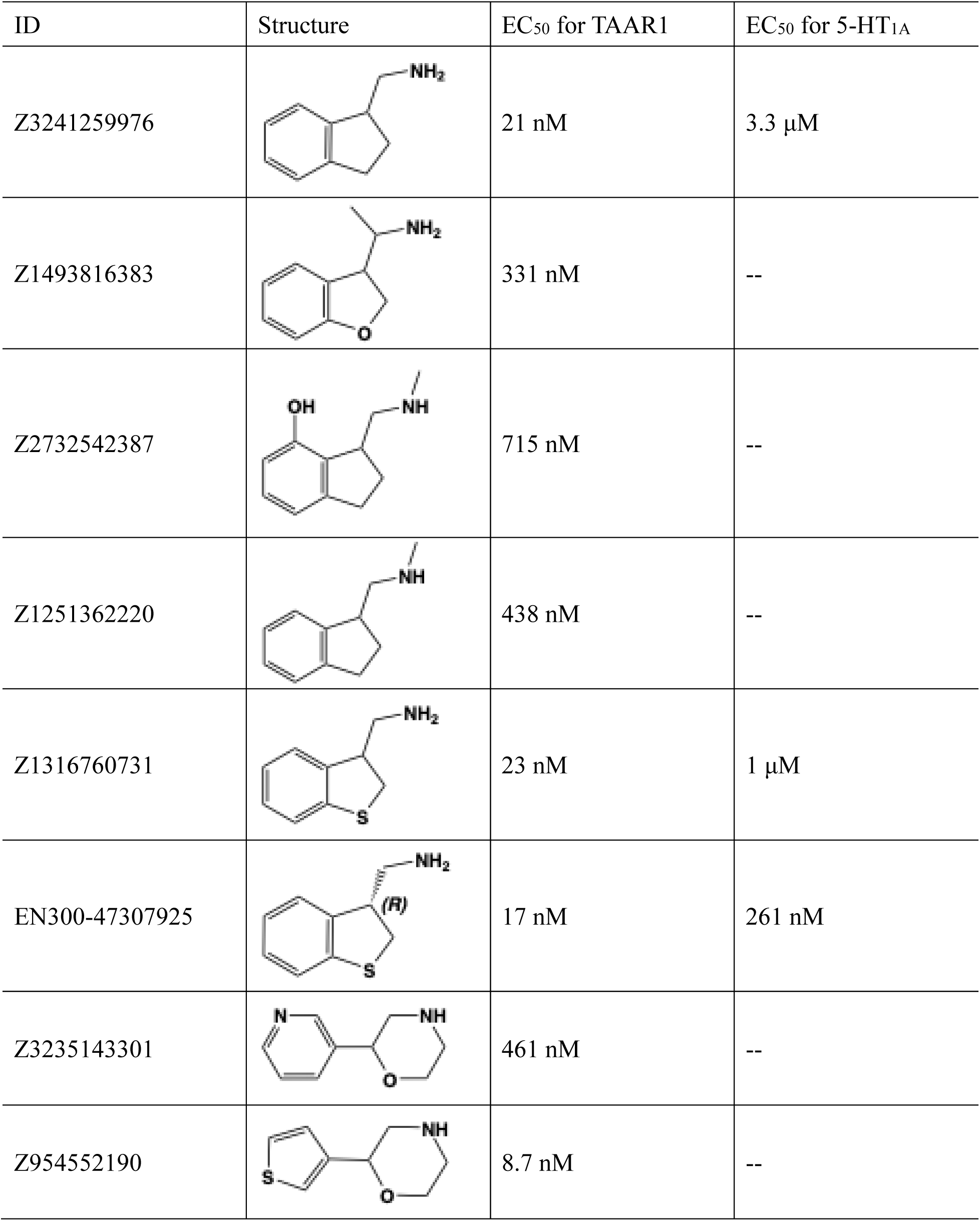
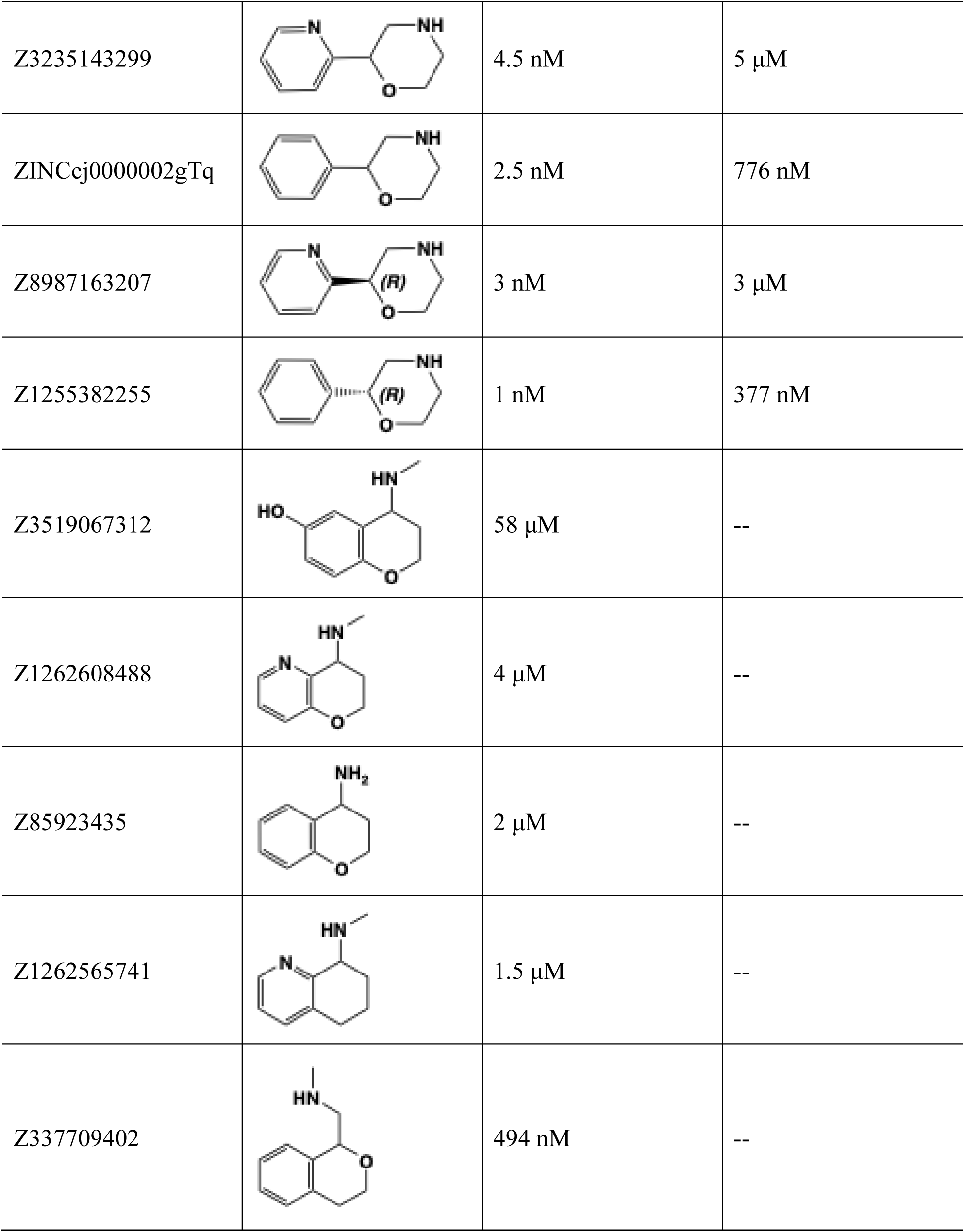
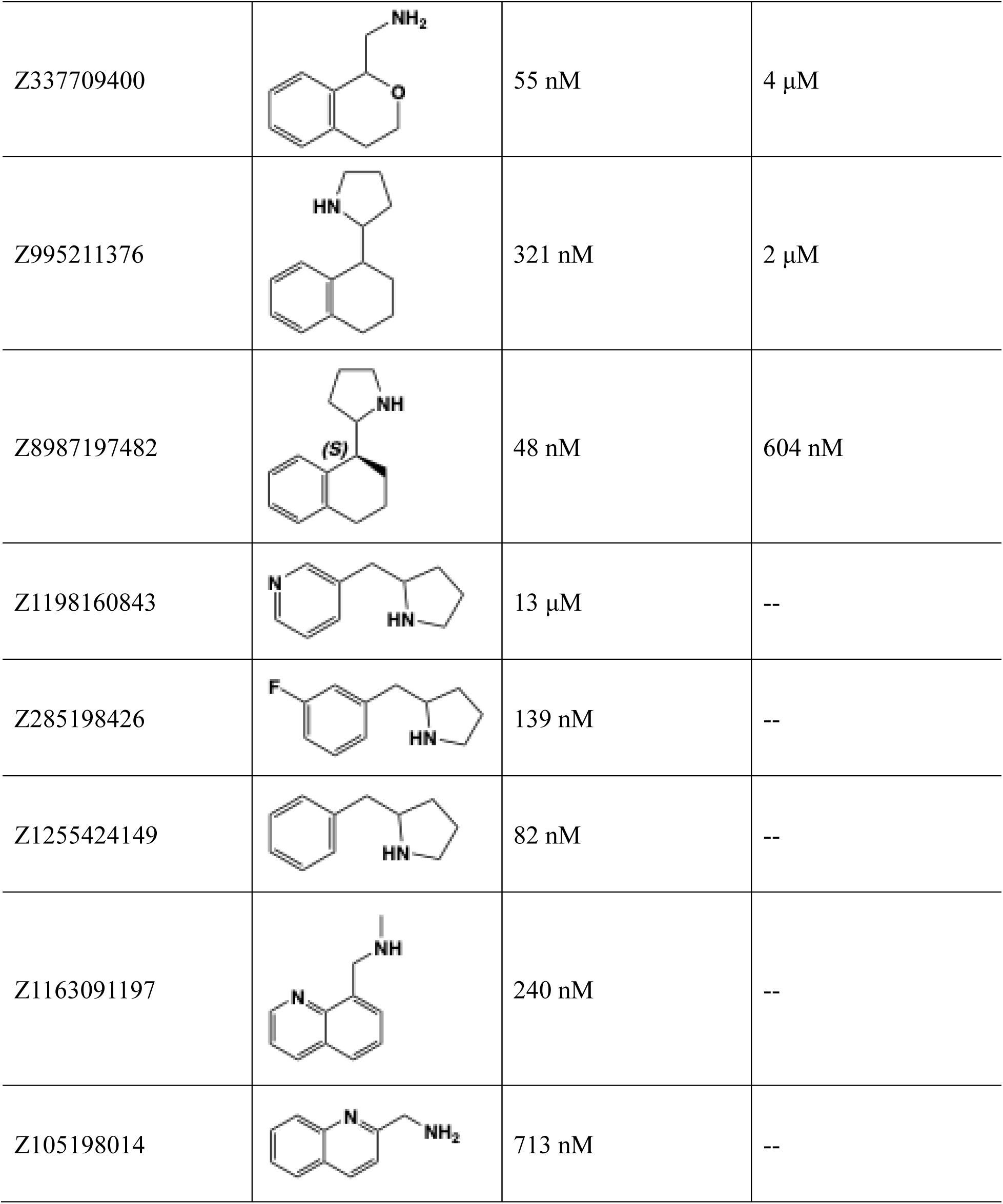
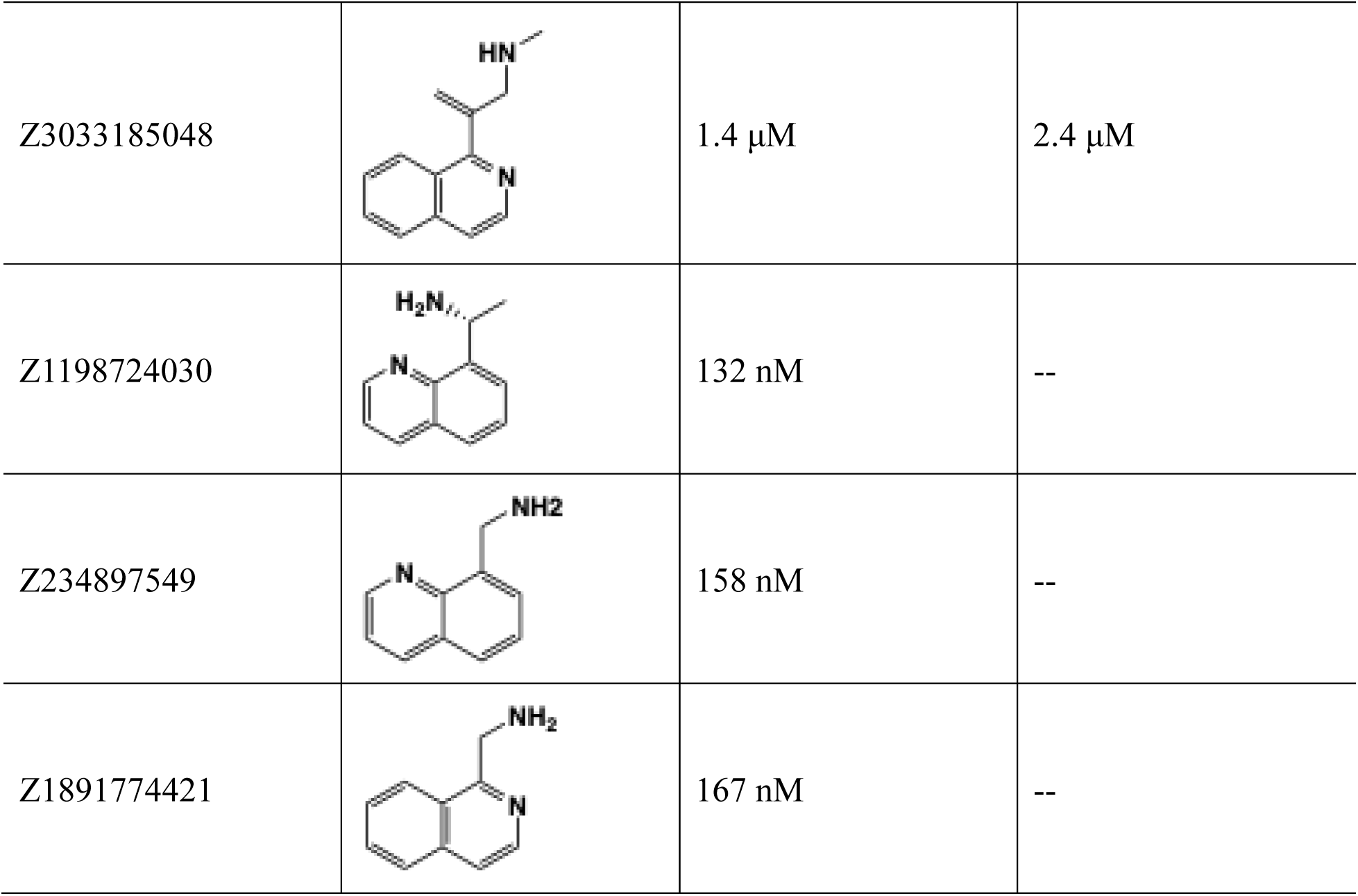
Optimized agonists from initial hits.

| ID | Structure | EC<sub>50</sub> for TAAR1 | EC<sub>50</sub> for 5-HT<sub>1A</sub> |
| --- | --- | --- | --- |
| Z3241259976 |  | 21 nM | 3.3 μM |
| Z1493816383 |  | 331 nM | -- |
| Z2732542387 |  | 715 nM | -- |
| Z1251362220 |  | 438 nM | -- |
| Z1316760731 |  | 23 nM | 1 μM |
| EN300-47307925 |  | 17 nM | 261 nM |
| Z3235143301 |  | 461 nM | -- |
| Z954552190 |  | 8.7 nM | -- |
| Z3235143299 | | 4.5 nM | 5 μM |
| ZINCcj0000002gTq |  | 2.5 nM | 776 nM |
| Z8987163207 | | 3 nM | 3 μM |
| Z1255382255 |  | 1 nM | 377 nM |
| Z3519067312 | | 58 μM | -- |
| Z1262608488 | | 4 μM | -- |
| Z85923435 | | 2 μM | -- |
| Z1262565741 | | 1.5 μM | -- |
| Z337709402 |  | 494 nM | -- |
| Z337709400 | | 55 nM | 4 μM |
| Z995211376 | | 321 nM | 2 μM |
| Z8987197482 |  | 48 nM | 604 nM |
| Z1198160843 | | 13 μM | -- |
| Z285198426 |  | 139 nM | -- |
| Z1255424149 |  | 82 nM | -- |
| Z1163091197 |  | 240 nM | -- |
| Z105198014 |  | 713 nM | -- |
| Z3033185048 | | 1.4 μM | 2.4 μM |
| Z1198724030 |  | 132 nM | -- |
| Z234897549 |  | 158 nM | -- |
| Z1891774421 |  | 167 nM | -- |

