## supplemental figure for "Structure based discovery of antipsychotic-like TAAR1 agonists"

### Supplementary Information

#### Supplementary Data

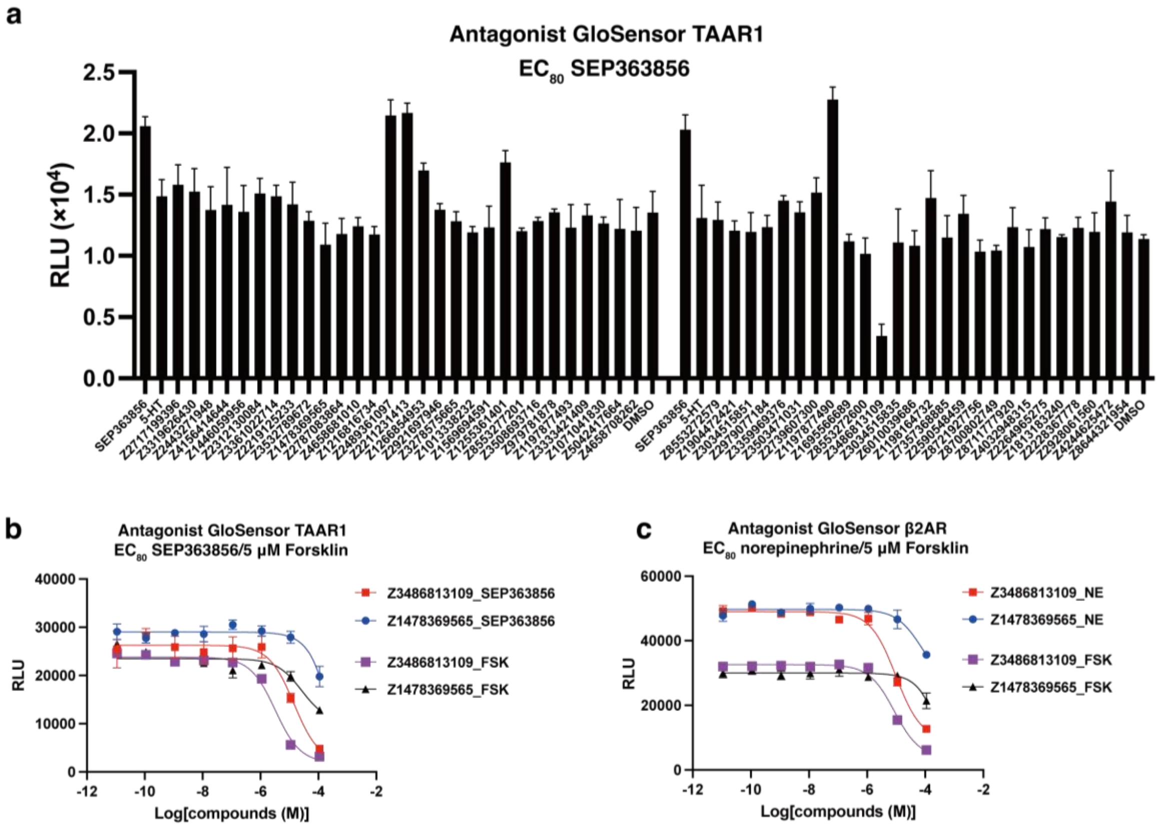


**Supplementary Fig. 1. Functional screening of predicted molecules against the TAAR1 in antagonist mode. a,** GloSensor responses of 55 predicted molecules at 10 µM in antagonist mode (cells were treated with $EC_{80}$ concentrations of agonist SEP363856). **b,** GloSensor assay in antagonist mode with cells expressing TAAR1 and treated with EC80 concentrations of agonist SEP363856 or 5 µM forskolin. **c,** GloSensor assay in antagonist mode with cells expressing β2AR and treated with EC80 concentrations of the agonist norepinephrine or 5 µM forskolin.

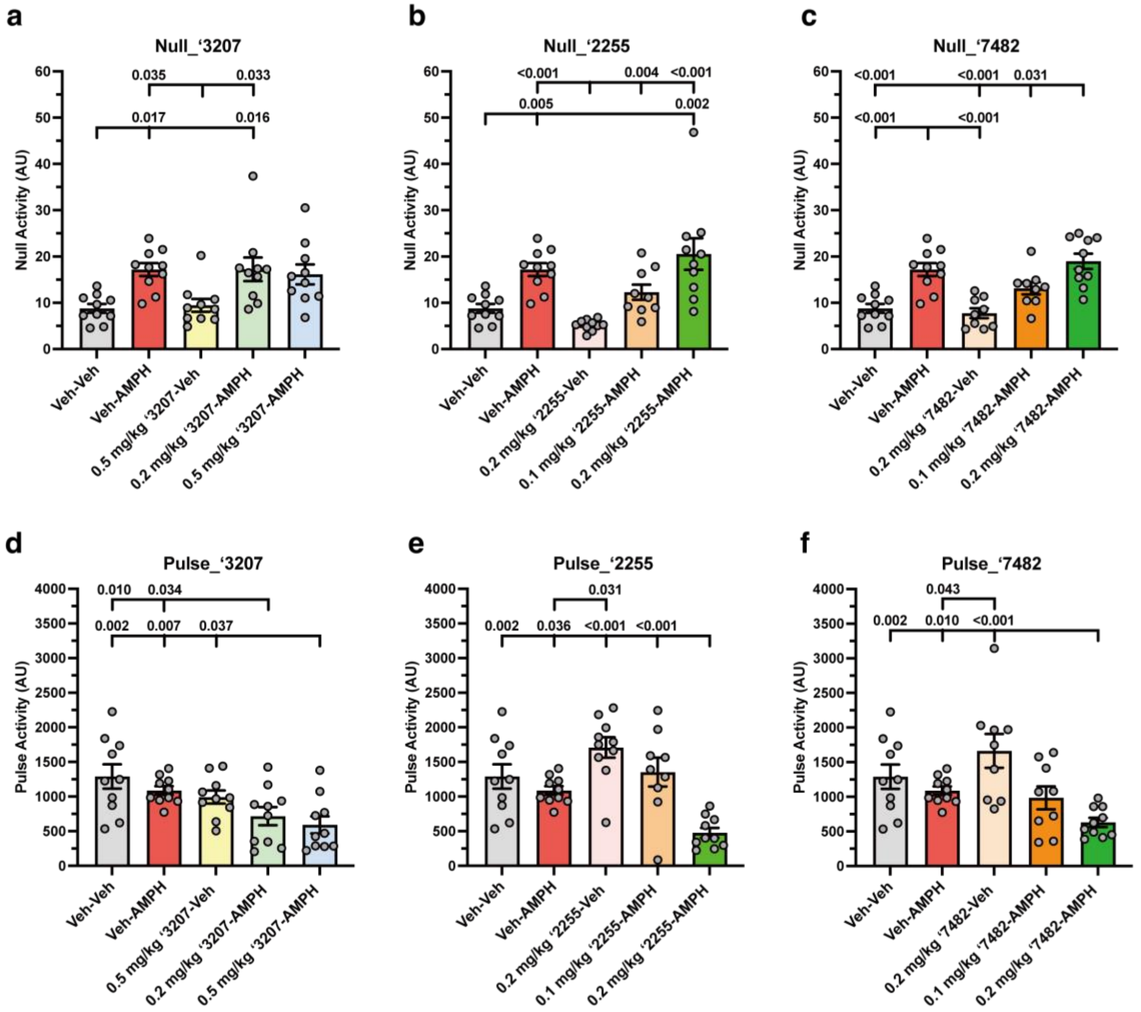


**Supplementary Fig. 2. Effects of ‘3207, ‘2255, and ‘7482 on null and pulse activities related to prepulse inhibition. a,** Effects of vehicle (Veh), 3 mg/kg amphetamine (AMPH), 0.5 mg/kg ‘3207, and ‘3207 with AMPH on null activities. **b,** Null responses to these treatments with ‘2255. **c,** Effects of these treatments with ‘7482 on null activities. **d,** Effects of Veh, AMPH, ‘3207, and ‘3207 with AMPH on pulse activities. **e,** Startle responses to these treatments with ‘2255. **f,** Effects of these treatments with ‘7482 on pulse activities. The data are presented as means and standard errors of the mean. In each panel the post-hoc p-value is provided for a specific comparison identified without a statistical value. The primary statistics are in Table S3.

**Supplementary Table 1. Cryo-EM data collection, refinement and validation statistics**

| | | |
|---|---|---|
| **Data Collection** | | |
| **Protein** | TAAR1- ‘7482 | TAAR1- ‘2255 |
| **Voltage (kV)** | 300 | 300 |
| **Detector** | Falcon 4 | Falcon 4 |
| **Pixel size (Å)** | 0.73 | 0.73 |
| **Defocus range (μm)** | 1.0-2.0 | 1.0-2.0 |
| **Electron dose ($e^-$/Å$^2$)** | 50 | 50 |
| **Frames per image** | 1.38 | 1.38 |
| **3D reconstruction** | | |
| **Final Particle number** | 137,014 | 72,449 |
| **Symmetry** | C1 | C1 |
| **Overall resolution (Å)** | 3.12 | 3.14 |
| **Model refinement** | | |
| **Model composition** | | |
| **Chains** | 6 | 6 |
| **Ligands** | 1 | 1 |
| **Non-hydrogen atoms** | 8231 | 8232 |
| **Protein residues** | 1040 | 1040 |
| **Bonds (RMSD)** | | |
| **Length (Å)** | 0.003 | 0.004 |
| **Angles (°)** | 0.567 | 0.571 |
| **Ramachandran plot (%)** | | |
| **Outliers** | 0.00 | 0.00 |
| **Allowed** | 2.24 | 2.63 |
| **Favored** | 97.76 | 97.37 |
| **Rotamer outliers (%)** | 0.11 | 0.00 |
| **MolProbity score** | 1.33 | 1.46 |
| **Clash score** | 5.21 | 6.07 |

**Supplementary Table 2. Pharmacokinetic properties of docking agonists.**

| **Compound** | **Sample** | **Administration** | **Dose, mg/kg** | **Pharmacokinetic Parameters** | | | | | |
|---|---|---|---|---|---|---|---|---|---|
| | | | | Tmax, min | Cmax, ng/ml | $AUC_{0 \to t\ min}$ (AUClast) ng*min/ml | $AUC_{0 \to \infty}$ (AUCINF_obs) ng*min/ml | $T_{1/2}$ (HL_Lambda_z), min | $K_{el}$ (Lambda_z), $min^{-1}$ |
| **'3027** | Plasma | IP | 10 | 30.0 | 6860 | 671000 | 673000 | 65.8 | 0.0105 |
| | Brain | | | 15.0 | 29800 | 2660000 | 2670000 | 66.9 | 0.0104 |
| | CSF | | | 15.0 | 3040 | 292000 | 295000 | 55.8 | 0.0124 |
| **'2255** | Plasma | IP | 10 | 15.0 | 3780 | 259000 | 260000 | 42.4 | 0.0164 |
| | Brain | | | 15.0 | 14600 | 882000 | 884000 | 38.9 | 0.0178 |
| | CSF | | | 15.0 | 1430 | 67400 | 75800 | 34.7 | 0.020 |
| **'7482** | Plasma | IP | 10 | 5.00 | 1810 | 134000 | 135000 | 65.0 | 0.0107 |
| | Brain | | | 30.0 | 11300 | 1870000 | 1880000 | 58.6 | 0.0118 |
| | CSF | | | 5.00 | 1750 | 123000 | 124000 | 64.0 | 0.0108 |

**Supplementary Table 3. Statistics for the prepulse inhibition studies.**

| Figure | Statistical Model | Variable | Degrees of Freedom | F- or H-statistic | *p*-value | # Mice per Group |
|---|---|---|---|---|---|---|
| Fig. 4a | RMANOVA[a] | PPI | 1.991,89.606 | 159.509 | <0.001 | 10 |
| | (PPI) | PPI x Treatment | 7.965,89.606 | 30.078 | 0.004 | |
| | | Treatment | 4,45 | 5.118 | 0.002 | |
| | Bonferroni | V-V *vs.* V-A[a] | | | 0.003 | |
| | (Treatment) | V-A *vs.* 0.5 Z3-V[a] | | | 0.050 | |
| | | V-A *vs.* 0.5 Z3-A[a] | | | 0.003 | |
| Fig. 4b | RMANOVA[a] | PPI | 1.997,87.876 | 246.416 | <0.001 | 9-10 |
| | (PPI) | PPI x Treatment | 7.989,87.876 | 3.270 | 0.003 | |
| | | Treatment | 4,44 | 7.190 | <0.001 | |
| | Bonferroni | V-V *vs.* V-A[a] | | | 0.002 | |
| | (Treatment) | V-V *vs.* 0.1Z2-A[a] | | | 0.009 | |
| | | V-A *vs.* 0.2Z2-V[a] | | | 0.014 | |
| | | V-A *vs.* 0.2Z2-A[a] | | | 0.009 | |
| | | 0.1Z2-A *vs.* 0.2Z2-A[a] | | | 0.038 | |
| | | 0.1Z2-A *vs.* 0.2Z2-A[a] | | | 0.038 | |
| Fig. 4c | RMANOVA[a] | PPI | 1.805,77.633 | 189.399 | <0.001 | 9-10 |
| | (PPI) | PPI x Treatment | 7.222,77.633 | 3.958 | <0.001 | |
| | | Treatment | 4,43 | 5.900 | <0.001 | |
| | Bonferroni | V-V *vs.* V-A[a] | | | 0.004 | |
| | (Treatment) | V-V *vs.* 0.1Z7-A[a] | | | 0.005 | |
| | | V-A *vs.* 0.2Z7-V[a] | | | 0.050 | |
| Suppl. Fig. S9a | One-Way ANOVA (Null Activity) | Treatment | 4,49 | 5.747 | <0.001 | 10 |
| | Bonferroni | V-V *vs.* V-A[a] | | | 0.017 | |
| | | V-V *vs.* 0.2Z3-A[a] | | | 0.016 | |
| | | 0.5Z3-V *vs.* V-A[a] | | | 0.035 | |
| | | 0.5Z3-V *vs.* 0.2Z3-A[a] | | | 0.033 | |
| Suppl. Fig. S9b | Kruskal-Wallis (Null Activity) | Treatment | 4,49 | 31.324 | <0.001 | 9-10 |
| | Dunn | V-V *vs.* V-A[a] | | | 0.005 | |

| | | | | | | |
|---|---|---|---|---|---|---|
| | | V-V *vs.* 0.2Z2-A[a] | | | 0.002 | |
| | | 0.2Z2-V *vs.* V-A[a] | | | <0.001 | |
| | | 0.2Z2-V *vs.* 0.1Z2-A[a] | | | 0.004 | |
| | | 0.2Z2-V *vs.* 0.2Z2-A[a] | | | <0.001 | |
| Suppl. Fig. S9c | One-Way ANOVA (Null Activity) | Treatment | 4,47 | 14.498 | <0.001 | 9-10 |
| | Bonferroni | V-A *vs.* V-V[a] | | | <0.001 | |
| | | V-A *vs.* 0.2Z7-V[a] | | | <0.001 | |
| | | 0.2Z7-A *vs.* V-V[a] | | | <0.001 | |
| | | 0.2Z7-A *vs.* 0.1Z7-V[a] | | | <0.001 | |
| | | 0.2Z7-A *vs.* 0.1Z7-A[a] | | | 0.031 | |
| Suppl. Fig. S9d | Kruskal-Wallis (Pulse Activity) | Treatment | 4,50 | 14.746 | 0.005 | 10 |
| | Dunn | 0.5Z3-A *vs.* V-V[a] | | | 0.002 | |
| | | 0.5Z3-A *vs.* V-A[a] | | | 0.007 | |
| | | 0.5Z3-A *vs.* 0.5Z3-V[a] | | | 0.037 | |
| | | 0.2Z3-A *vs.* V-V[a] | | | 0.010 | |
| | | 0.2Z3-A *vs.* V-A[a] | | | 0.034 | |
| Suppl. Fig. S9e | One-Way ANOVA (Pulse Activity) | Treatment | 4,48 | 10.434 | <0.001 | 9-10 |
| | Bonferroni | 0.2Z2-A *vs.* V-V[a] | | | 0.002 | |
| | | 0.2Z2-A *vs.* V-A[a] | | | 0.036 | |
| | | 0.2Z2-A *vs.* 0.2Z2-V[a] | | | <0.001 | |
| | | 0.2Z2-A *vs.* 0.1Z2-A[a] | | | <0.001 | |
| | | V-A *vs.* 0.2Z2-V[a] | | | 0.031 | |
| Suppl. Fig. S9f | Kruskal-Wallis (Pulse Activity) | Treatment | 4,48 | 17.340 | 0.002 | 9-10 |
| | Dunn | 0.2Z7-A *vs.* V-V[a] | | | 0.002 | |
| | | 0.2Z7-A *vs.* V-A[A] | | | 0.010 | |
| | | 0.2Z7-A *vs.* 0.2Z7-V[A] | | | <0.001 | |
| | | 0.2Z2-v *vs.* 0.1Z7-A[A] | | | 0.043 | |

[a]Abbreviations: RMANOVA, repeated measures ANOVA; PPI, prepulse inhibition; V-V, Vehicle-Vehicle; V-A, Vehicle-3 mg/kg Amphetamine; 0.5Z3-V, 0.5 mg/kg Z3207-Vehicle; 0.2Z3-A, 0.2 mg/kg Z3207-3 mg/kg Amphetamine; 0.5Z3-A, 0.5 mg/kg Z3207-3 mg/kg Amphetamine; 0.2Z2-

V, 0.2 mg/kg Z2255-Vehicle; 0.1Z2-A, 0.1 mg/kg Z2255-3 mg/kg Amphetamine; 0.2Z2-A, 0.2 mg/kg Z2255-3 mg/kg Amphetamine; 0.2Z7-V, 0.2 mg/kg Z7482-Vehicle; 0.1Z7-A, 0.1 mg/kg Z7482-3 mg/kg Amphetamine; 0.2Z7-A, 0.2 mg/kg Z7482-3 mg/kg Amphetamine.
